## Supplemental Methods and Figures for "Community assessment of methods to deconvolve cellular composition from bulk gene expression"

### Supplementary Methods and Figures for Community assessment of methods to deconvolve cellular composition from bulk gene expression


**Supplemental Methods** **2**
- Figure and table generation 2
- Aginome-XMU deconvolution method 6
  - Summary Sentence 6
  - Introduction 6
  - Methods 7
    - sub-Challenge 1 7
      - Prediction method 7
      - Prediction output 7
      - Normalization/pre-processing data 8
      - Training models 8
    - sub-Challenge 2 9
      - Results 9
  - Conclusion/Discussion 11
- DA_505 deconvolution method 11
  - Summary Sentence 11
  - Background/Intro 12
  - Methods 12
    - Normalization/pre-processing data 13
    - Prediction method 15
    - Prediction output 16
  - Conclusion/Discussion 17
  - Authors Statement 18
- mitten_TDC19 deconvolution method 18
  - Summary Sentence 18
  - Background/Intro 18
  - Methods 18
  - Conclusion/Discussion 19
  - Authors Statement 19
- Biogem deconvolution method 19
  - Summary Sentence 19
  - Background/Intro 19
  - Methods 20

Data collection and preprocessing 20
Feature selection 22
Signature matrix 22
Prediction method 22
Prediction output 23
Variations in 2nd and 3rd submission 23
Variations between sub-Challenge 1 (coarse-grained sub-Challenge) and sub-Challenge 2 (fine-grained sub-Challenge) 25
Conclusion/Discussion 25
Authors Statement 26
IZI deconvolution method 26
**Supplemental Figures** **28**
**References** **48**
**Figure Legends** **53**

### Supplemental Methods

#### Figure and table generation

The following scripts in the github repository
https://github.com/Sage-Bionetworks/Tumor-Deconvolution-Challenge
were used to generate the indicated tables and figures:

| Table number | Table name | Script that produced the table |
|---|---|---|
| S1 | analysis/validation-analysis/figs/rerun-validation-mixture-and-distribution-effects.tsv | analysis/validation-analysis/plot-bootstrap-rerun-validation.R |
| S2 | analysis/sample-level-analysis/figs/figs/sample-level-comparison.tsv | analysis/sample-level-analysis/plot-sample-level-summaries.R |
| S10 | training-sample-metadata/geo-expression-array-immune-cells.csv | training-sample-metadata/download-and-format-training-sample-metadata.R |
| S11 | training-sample-metadata/geo-rnaseq-immune-cells.csv | training-sample-metadata/download-and-format-training-sample-metadata.R |

| | | |
|---|---|---|
| 1 | method-annotation-table/deconv-method-description-table-1.pdf | method-annotation-table/deconv-method-description-table-1.tex |
| 2 | method-annotation-table/deconv-method-description-table-2.pdf | method-annotation-table/deconv-method-description-table-2.tex |
| S12 | analysis/timing/method-run-times.tsv | analysis/timing/format-timing-results.R |
| S13 | training-sample-metadata/SampleDetails.kallisto.csv.gz | training-sample-metadata/download-and-format-training-sample-metadata.R |
| S14 | training-sample-metadata/DA505_training_metadata.csv | https://figshare.com/s/944b97b01c91763e8dcd |
| S15 | training-sample-metadata/detailedMetadataOfSamples.xlsx | training-sample-metadata/download-and-format-training-sample-metadata.R |
| S16 | training-sample-metadata/Metadata.csv | training-sample-metadata/download-and-format-training-sample-metadata.R |
| S19 | analysis/cancer-validation/figs/cancer-validation-dataset-comparison-pvals.tsv | analysis/cancer-validation/score-cancer-datasets.R |

| Figure number | File name | Script that produced the figure |
|---|---|---|
| Fig S1 | analysis/summary/purified-samples-marker-heatmap-protein-coding-genes-no-title.png | analysis/summary/plot-marker-heatmap.R |
| Fig 2 | analysis/validation-analysis/figs/fig-validation-round-1-performance.png | analysis/validation-analysis/perform-bootstrap-rerun-validation.R, plot-bootstrap-rerun-validation.R |

| | | |
|---|---|---|
| Fig S2 | external-figs/da_505-supp-fig 1.png | DA505 writeup: https://www.synapse.org/#!Synapse:syn20674744/wiki/603943 |
| Fig S3 | external-figs/Biogem_ComparisonDataset.png | Biogem writeup: https://www.synapse.org/#!Synapse:syn20551146/wiki/594139 |
| Fig S4 | analysis/validation-analysis/figs/fig-validation-performance-across-rounds.png | make-validation-performance-figs.R |
| Fig S5 | analysis/validation-analysis/figs/fig-validation-all-performance.png | analysis/validation-analysis/plot-bootstrap-rerun-validation.R |
| Fig 3 | analysis/validation-analysis/figs/fig-validation-round-1-strip-and-heatmap-merged-cell-type.png | analysis/validation-analysis/plot-bootstrap-rerun-validation.R |
| Fig S6 | analysis/validation-analysis/figs/fig-validation-round-1-merged-strip-cell-type.png | analysis/validation-analysis/plot-bootstrap-rerun-validation.R |
| Fig S7 | analysis/validation-analysis/figs/fig-validation-heatmap-round-1-coarse-and-fine-cell-type.png | analysis/validation-analysis/plot-bootstrap-rerun-validation.R |
| Fig S8 | analysis/validation-analysis/figs/fig-validation-round-1-coarse-and-fine-strip-cell-type.png | analysis/validation-analysis/plot-bootstrap-rerun-validation.R |
| Fig S9 | analysis/validation-analysis/figs/fig-validation-heatmap-rounds-2-and-3-merged-cell-type.png | analysis/validation-analysis/plot-bootstrap-rerun-validation.R |
| Fig S10 | analysis/validation-analysis/figs/fig-validation-round-2-merged-strip-cell-type.png | analysis/validation-analysis/plot-bootstrap-rerun-validation.R |

| | | |
|---|---|---|
| Fig S11 | analysis/validation-analysis/figs/fig-validation-heatmap-round-2-coarse-and-fine-cell-type.png | analysis/validation-analysis/plot-bootstrap-rerun-validation.R |
| Fig S12 | analysis/validation-analysis/figs/fig-validation-round-2-coarse-and-fine-strip-cell-type.png | analysis/validation-analysis/plot-bootstrap-rerun-validation.R |
| Fig S13 | analysis/validation-analysis/figs/fig-validation-round-3-merged-strip-cell-type.png | analysis/validation-analysis/plot-bootstrap-rerun-validation.R |
| Fig S14 | analysis/validation-analysis/figs/fig-validation-heatmap-round-3-coarse-and-fine-cell-type.png | analysis/validation-analysis/plot-bootstrap-rerun-validation.R |
| Fig S15 | analysis/validation-analysis/figs/fig-validation-round-3-coarse-and-fine-strip-cell-type.png | analysis/validation-analysis/plot-bootstrap-rerun-validation.R |
| Fig 4 | analysis/sample-level-analysis/figs/sample-level-metric-swarm-round-1.png | analysis/sample-level-analysis/plot-sample-level-summaries.R |
| Fig 5 | analysis/specificity-analysis/figs/spillover-summary.png | analysis/specificity-analysis/plot-validation-spillover-results.R |
| Fig S16 | analysis/specificity-analysis/figs/spillover-all-scores-coarse-grained-round-1.png | analysis/specificity-analysis/plot-validation-spillover-results.R |
| Fig S17 | analysis/specificity-analysis/figs/spillover-all-scores-fine-grained-round-1.png | analysis/specificity-analysis/plot-validation-spillover-results.R |
| Fig 6 | analysis/in-silico-admixtures/figs/sensitivity-spikein-and-summary.png | analysis/in-silico-admixtures/plot-sensitivity-results.R |
| Fig S18 | analysis/cancer-validation/figs/fig-cancer-validation-heatm | analysis/cancer-validation/score-cancer-datasets.R |

| | | |
|---|---|---|
| | ap-wu-and-pelka.png | |
| Fig S19 | analysis/cancer-validation/figs/fig-cancer-validation-heatmap-all.png | analysis/cancer-validation/score-cancer-datasets.R |
| Fig 7 | analysis/cancer-validation/figs/fig-cancer-validation-per-cell-type.png | analysis/cancer-validation/score-cancer-datasets.R |
| Fig S20 | analysis/cancer-validation/figs/fig-cancer-validation-legend-edited.png | analysis/cancer-validation/score-cancer-datasets.R |
| Fig S21 | analysis/summary-table/figs/binned-score-heatmap-score-sorted-ignore-nas-comparator.png | analysis/summary-table/make-summary-table.R |

### Aginome-XMU deconvolution method

#### Summary Sentence

We used a deep learning-based prediction model for cell fraction prediction from bulk RNA-seq data.

#### Introduction

Numerous computational deconvolution methods have been proposed to estimate the abundance of individual cell types from bulk RNA-seq data of heterogeneous tissues. Unfortunately, there are still many difficulties hindering the performance of these algorithms, such as the collinearity of expression from different cell types, lack of specific marker genes, and the difficulty in dealing with batch effects of technical variation of data from different platforms.

In recent years, we have witnessed a paradigm shift in the machine learning community. In contrast to traditional feature engineering approaches where feature extraction and classification are optimized separately, new techniques based on learning internal representations from data directly have been explored. These data-driven approaches, especially deep learning, have allowed significant research breakthroughs and have rapidly spread across multiple application domains such as computer vision, audio recognition, and human language processing.

In this work, we hypothesized that by using deep learning, we could capture the intrinsic relationship between cellular composition of tissue and its bulky gene expression profile without the need for manual feature selection or marker gene identification. To test this hypothesis, we developed a deep learning-based cell type fraction prediction model with in silico mixing training data from multiple microarray and RNA-seq (Table S13), as well as scRNA-seq, datasets. We tested its performance on our testing sets comprised of nine public datasets (GSE64385, GSE65133, GSE106898, GSE107011, GSE64655, GSE127813, SDY67, GSE59654, GSE107990) and datasets from three public leaderboard rounds. We found that the algorithm shows better accuracy and stability in comparison with existing methods on our testing sets.

#### Methods

##### sub-Challenge 1

###### Prediction method

We used a deep learning-based prediction model for cell fraction prediction from bulk RNA-seq data. Briefly, we trained deep feed-forward, fully connected neural networks (multilayer perceptron networks) on in silico mixed training data. The network consists of one input layer, five fully connected hidden layers and one output layer and was implemented with the PyTorch framework (v1.0.1) in Python (v3.7.3). The detailed description of this deep learning-based model have been published.[1]

In addition to the deep learning-based method, in our submission we use an ensemble method to further improve the prediction results. Specifically, we noticed that summarization method (MCP-counter[2]) performs better on neutrophils, fibroblasts, and endothelial cells on our testing sets. Therefore, MCP-counter was used to predict the cell type proportion in our second submission to sub-Challenge 1.

##### Prediction output

Our deep learning-based method outputs the absolute fraction of target cell types. Hence, the results can be compared across both cell types and samples. The method can score all cell types present in the sample including immune, stroma and cancer. Non-negative or sum to one constraint were currently not enforced in our model. As the deep learning model produces absolute cell fraction prediction, the fraction of other cell types can be calculated by subtracting the total of the predicted fractions of all the supported cell types from 1.

##### Normalization/pre-processing data

We use min-max scaling to normalize the expression data. In the deep-learning model, we do not apply feature selection. Instead, all the available gene expression data, after intersection of the genes of the training data with the genes of the test data, were used as our input to the deep learning model.

##### Training models

We mixed the purified expressions of 8 cell types in random proportions, and at the same time mixed a part of the unknown cell lines (e.g., cancer cell line). These purified expression samples came from microarray, RNA-seq, scRNA-seq. The expression of a simulated sample $\boldsymbol{e}$ was calculated as

$$\boldsymbol{e} = \sum_{k=1}^{C} p_k \boldsymbol{\epsilon}_k,$$

where $C$ is the number of cell types involved in mixing, $p_k$, $k = 1, \ldots, C$ are random variables with Dirichlet distribution that determined the fractions of different cells in the in silico mixed sample, and $\boldsymbol{\epsilon}_k$ is the expression profile of a randomly selected purified sample of cell type $k$ from the respective RNA-seq or microarray dataset.

For scRNA-seq dataset, $\boldsymbol{e}$ is given by

$$e = \sum_{k=1}^{C} \sum_{j=1}^{n_k} \epsilon_{kj},$$
,

where $n_k = 500 \cdot p_k$ is the number of cells of type $k$ extracted randomly from scRNA-seq datasets for mixing, and $\epsilon_{kj}$ denote their expression profiles. Note that in this case $e$ were further TPM-normalized before used as training data.

In this way, the expression of heterogeneous samples in the real-world tumor microenvironment is simulated, and large number of samples can be generated as training data.

In addition, we also used a previously published[1] in silico mixing method to expand a large number of training samples through mixing RNA-seq dataset SDY67 with ground truth and scRNA-seq data.

We used bagging to select 30%-100% training datasets for model training. In model training, we used cross-validation through selecting 20% training data as a validation set to evaluate the training degree of the model. We used early stopping such that training is stopped when the loss on the validation set is not reduced after 10 epochs.

##### sub-Challenge 2

We used a similar approach as in sub-Challenge 1 to create our prediction model. The only exception is that in our third submission, xCell[3] was used to create prediction of cell type myeloid dendritic cells as it performs better than other methods on this cell type on our testing sets.

###### Results

In the coarse cell type track, we made two submissions. In the first submission, we use a deep learning-based prediction model as described above on all cell types. For each cell type, we chose the best model that achieved the highest Pearson correlation results on our testing datasets. The grand mean of the Pearson correlation over all cell types was 88.3%.

We noticed that on our testing datasets, the deep learning-based model did not outperform marker gene-based models such as MCP-counter on some cell types, namely, neutrophils, fibroblasts, and endothelial cells. We believe that these cell types have highly specific marker genes, therefore it is relatively easy to predict using marker gene-based models. Therefore, we replaced the method with MCP-counter on these three cell types to profile the abundance of the cells in the second submission. The grand mean of the Pearson correlation over all cell types increased to 92% in this submission. The results from our two coarse-grained submissions are as follows:

| Team | Aginome-XMU | Aginome-XMU |
|---|---|---|
| objectId | 9704292 | 9704388 |
| submission | first submission | second submission |
| B.cells | 0.894/DNN | 0.894/DNN |
| CD4.T.cells | 0.862/DNN | 0.862/DNN |
| CD8.T.cells | 0.896/DNN | 0.896/DNN |
| monocytic.lineage | 0.931/DNN | 0.931/DNN |
| NK.cells | 0.826/DNN | 0.826/DNN |
| neutrophils | 0.94/DNN | 0.993/MCP-counter |
| fibroblasts | 0.851/DNN | 0.98/MCP-counter |
| endothelial.cells | 0.864/DNN | 0.98/MCP-counter |
| Grand mean | 0.883 | 0.92 |

In the fine cell type track, we made three submissions. In the first submission, we used a deep learning model to predict the fraction of 13 fine cell types. The grand mean of the Pearson correlation was 77.9%. To further improve the performance, we replaced deep learning-based model with MCP-counter for three cell types (neutrophils, fibroblasts, and endothelial cells) on which marker gene-based model performs better than deep learning-based model in the testing set, and obtained a grand mean of 78.6%. In the final submission, we further replaced the model for myeloid dendritic cells to xCell, which performed very well on this cell type in our test data. The final grand mean score was 81.9%. The results from our three fine-grained submissions are as follows:

| Team | Aginome-XMU | Aginome-XMU | Aginome-XMU |
|---|---|---|---|
| objectId | 9704261 | 9704399 | 9704690 |
| submission | first submission | second submission | third submission |
| naive.B.cells | 0.939/DNN | 0.939/DNN | 0.939/DNN |
| naive.CD4.T.cells | 0.771/DNN | 0.771/DNN | 0.771/DNN |
| memory.CD4.T.cells | 0.523/DNN | 0.523/DNN | 0.523/DNN |
| regulatory.T.cells | 0.8/DNN | 0.8/DNN | 0.8/DNN |
| naive.CD8.T.cells | 0.875/DNN | 0.868/DNN | 0.875/DNN |
| memory.CD8.T.cells | 0.645/DNN | 0.573/DNN | 0.645/DNN |
| NK.cells | 0.91/DNN | 0.91/DNN | 0.91/DNN |
| monocytes | 0.733/DNN | 0.733/DNN | 0.733/DNN |
| myeloid.dendritic.cells | 0.468/DNN | 0.468/DNN | 0.815/xCell |
| macrophages | 0.672/DNN | 0.672/DNN | 0.672/DNN |
| neutrophils | 0.915/DNN | 0.999/MCP-counter | 0.999/MCP-counter |
| fibroblasts | 0.928/DNN | 0.981/MCP-counter | 0.981/MCP-counter |
| endothelial.cells | 0.95/DNN | 0.979/MCP-counter | 0.979/MCP-counter |
| Grand mean | 0.779 | 0.786 | 0.819 |

#### Conclusion/Discussion

Looking forward, we believe that the following points can be further elaborated to improve the cell fraction deconvolution performance.

First, we used all the gene expression data for genes available in both training and testing datasets as the input of our prediction model. However, further analysis showed that different genes contribute differently to the prediction results. Therefore, it is possible to significantly reduce the number of genes used as prediction input by identifying a small subset of genes that contribute most to the prediction results.

Secondly, we found that although the deep learning-based model performs relatively well on all cell types, for some cell types, shallow models such as MCP-counter or xCell perform better. Therefore, it seems that both deep and shallow models provide complementary information to some extent, and a late fusion strategy to combine both models could potentially be used to further improve the performance. Note that although shallow models such as MCP-counter or xCell perform better on those cell types, they do not produce absolute cell fractions as the deep learning-based model does. Hence, the late fusion strategy could potentially bring additional advantage to unify different kinds of output scores to absolute scores that can be compared across cell types and samples.

Finally, as suggested in reported results,[1] it may be challenging to produce a one-size-fits-all prediction model that consistently performs well across datasets of different sources due to the existence of batch effects and technical variations from different experiment sites. A better solution seems to be training a dataset-specific prediction model. Preliminary results on datasets with a small number of calibration samples with ground truth cell fractions are very promising in our experiments.

### DA_505 deconvolution method

#### Summary Sentence

Our method utilizes Random Forest regression to select the most significant features associated with cell-type proportions and (support vector or penalized linear) regression to predict different cell type proportions in complex RNA admixtures using an approach to normalize and mix samples of purified cell types.

#### Background/Intro

Usually, RNA-Seq gene expression analysis is approached by applying a plethora of scaling and normalization methods to the sample's raw counts to apply different data analysis methods on the processed estimates. Although current techniques have proven to be solid and robust, many rely on inter-sample information gathering to perform their normalization procedures such as quantile normalization or using the library size to scale the expression counts (Voom normalization). This presents great obstacles when these methods are applied to samples that share the same biological origin but are from different technical contexts. These issues represent a great obstacle not only when we want to train a model using multiple datasets from different research centers, but also when we wish to apply our prediction models to new unseen samples from different centers or as isolated cases (i.e., when we desire to apply the model as a product to aid medical diagnosis and decide treatment).

Here we propose a normalization method to adjust count values measured on different scales into a common scale disregarding the sample counts' prior probability distribution or library size, utilizing a transformed ranking system under the assumption that the features expression order in a given sample would be unaltered under different scales and probability distributions. In particular, this approach allowed us to mix RNA-Seq experiments of purified cell lines into complex admixtures to train different machine learning models to predict cell-type proportions in new unseen cases.

To test our hypothesis we trained three mainstream Machine Learning models for each cell type using the admixtures created from 23 different datasets of bulk RNA-Seq experiments of purified cell types. Partial Least Squares (PLS), Penalized linear regression (GLMNET), and Support Vector Regressors (SVR) were used combined with a feature selection approach using Random Forest (RF) to reduce the dimensionality of the problem.

#### Methods

In both sub-Challenge 1 and sub-Challenge 2 the same approach was used with a variation in the creation of the admixtures used to train the models. In both cases, our approach performed well, which shows the versatility of the framework proposed.

Intuitively, the workflow can be divided into three phases: 1) Data curation, downloading, and pre-processing. 2) Data normalization, quality control and admixture creation, and 3) model training, hyperparameter tuning, and model selection. Since the SVR was the model with the best results, we will describe only that model here. The SVR model was used for our third submissions, whereas GLMNET was used for our first two submissions. The whole workflow is depicted in Fig. S2 and each step will be explained in the subsequent sections.

##### Normalization/pre-processing data

We believe that the most important step in our framework was the data pre-processing and normalization.

To train our model 23 different sequencing datasets were manually curated and downloaded from public repositories where purified immune, stromal, and cancer cell types from different origins and conditions were sequenced using different sequencing platforms (Genome Analyzer II, NextSeq500, Hiseq2000, 2500, 4000, Novaseq600, etc). The selection criteria to choose the different experiments were:

- Human isolated stromal, immune or cancer cell types
- No molecular modifications of the cells were performed on the samples
- The samples represent biologically plausible conditions

Once downloaded, samples were manually labeled in a unified coarse-grain or fine-grain classification. For the coarse-grain class the subtypes cancer, fibroblast, endothelial, NK cells, Neutrophils, monocytic lineage, B cells, CD4 T cells, CD8 T cells and Adipocytes were considered. Other cell types present in the used datasets were classified as others. For the fine-grain labels, immune cells were further categorized. Monocytic lineage was divided into monocytes, macrophages and myeloid dendritic cells. B, CD4, and CD8 cells were further classified into naïve and memory, and the regulatory T cell label was also included for CD4 cells. Also, the others label was used.

The metadata of the datasets is included in Table S14.

Once all the samples were downloaded, a quality control analysis was performed upon all available samples using FASTQC software.[4]

After performing the first fastqc analysis we noticed that some sample sets, e.g. ERP002049, had samples with reads of very poor quality which lead us to trial a series of trimmomatic settings to filter out poor quality reads.[5] Our strategy was to seek a balance between achieving the highest quality and retaining the most number of fragments possible for each sample. As the alignment software used (Kallisto) only looks for exact matches,[6] we placed strong value on high quality reads to reduce the possibility of misalignments. By default Kallisto uses 31 bp kmers to align the reads to features, considering this we opted for a MINLEN setting of 36 and used the trimmomatic MAXINFO option to perform an adaptive quality trim, balancing the benefits of retaining longer reads against the costs of retaining bases with errors. Using these settings mean fragment lengths for each sample set typically ranged from 40 - 49 bp. Eight samples from SRP031776 had read lengths of 27bp before trimming and were omitted from our analysis.

Kallisto pseudoalignment requires the strandedness of the library to be provided, which we verified prior to transcript count quantification using Kallisto's default parameters. The transcript – count matrix was then collapsed to a gene – count matrix using the tximport R package.[7]

Once the count matrix was obtained, all samples were normalized using an ad-hoc developed normalization procedure we named RHINO. The rationale behind this approach is based on the observation of methods that use library size to scale counts, such as Voom transformation,[8] usually perform satisfactorily when comparing samples from the same origin, but fail to correct probability distribution differences between samples from different experimental contexts. Due to this observation, other widely used methods (i.e. quantile normalization) rely on taking all the samples to a single probability distribution under the assumption that all the sample counts come from the same underlying distribution. This second approach usually corrects satisfactorily the inequalities found from different sample centers, nevertheless, it fails to correct for systematic overdispersion of the counts from low expression genes (own observations, manuscript under preparation).

Taking these observations into consideration, we propose a new transformation method to normalize sample counts across datasets of different origins;

$$X\prime = -log_2\left[rank\left(-1X\right)\right] + log_2\left(n\right)$$

Where $X$ is the count matrix of a given sample with dimensions $n * 1$, with $n$ being the number of features in the count matrix. With this method, we only take into account the ranks of the expressed features using the max form to account for ties in the rankings.

Once samples were collected and normalized, a sample selection procedure was performed to discard possible low quality or mislabeled purified samples. Briefly, a penalized multinomial classifier was trained with the complete dataset. Then, we selected the non-zero coefficients of the model and used this subset of features to calculate Pearson's distance between samples.

Since some cell types might be similar or related in their origin (for example CD4 and CD8 T cells), we used a community detection clustering approach to detect communities of samples from the same origin. In particular, samples with Pearson's correlation coefficients greater than 0.8 were considered to be similar, information that was used to perform a Louvain clustering procedure that detected clearly defined communities of unique cell populations.[9] Samples belonging to these clearly defined clusters were used to create the admixtures while samples not falling into these groups were discarded.

Once the prototypic samples for each cell type were defined, 75% of the samples of each cell population were used in a training set, and the remaining 25% of the samples were used as a validation set to perform parameter tuning.

To create the admixtures to perform the model training, a data augmentation approach was used where samples from within the same cell type population were mixed to create new purified samples, and these new samples were mixed in silico with samples from other cell populations in known proportions to create a training and validation admixture set of 2000 and 500 samples respectively.

Using these two sets a Random Forest (RF) for feature selection was performed.[10]

The RF algorithm consists of a collection of tree-structured regressors. In general, each tree grows with respect to a random subset of the input dataset independent and identically distributed. The strategy known as Random Input Selection (RIS) is used to generate different trees. The algorithm chooses randomly a subset S with M features from the original set of n features and seeks within S the best feature to split a node of the tree according to some purity measure. Therefore a regression tree is found with M feature subset for each subset S. The final output for a given input data point is calculated using the average of all the regression trees predicted values. The parameters considered during the tuning process of the algorithm are the number of trees (ntree) and the number of features considered on each tree (mrty). The parameter ntree was fixed in 500 while the mrty was exponentially ranged between 2 and 200.

By using RF it is possible to obtain a rank of the most important features used for the deconvolution problem. By default, RF uses the mean decrease impurity and is related to the total decrease in node impurity averaged over all trees The use of RF for feature selection over linear regressors coefficients is preferred because of its capability of dealing with non-linear regression as well as a simple straightforward approach for measuring feature importance.

##### Prediction method

Once we obtained the training and validation In Silico admixtures we decided to use three classical regression methods to predict the different cell type proportions under the assumption that some features (genes) alone or in combination can act as specific cell subtype biomarkers that can be detected over biologically common or unspecific gene expression patterns. The three methods tested were Support Vector Regression (SVR), Partial Least Squares (PLS), and

Penalized Linear regression (GLMNET) but other methods could also be applied. In particular, for each cell line, we trained a model that was fine-tuned using a validation set made out of a set of left out samples for this purpose and the decision of the best performing method was made by the comparison of the mean Pearson's correlation coefficient over all cell lines prediction in the test sets provided in the first three phases of the challenge.

Here we will further explain the SVR approach since it was the best performing submitted model in the final submission (and used for Round 3). We used a GLMNET approach in Rounds 1 and 2.

The results of our tested models for the coarse-grained sub-Challenge are here (https://rpubs.com/harpomaxx/devcon2020-5-newmix) and for the fine-grained sub-Challenge here (https://rpubs.com/harpomaxx/devcon2020-finegrain).

Support Vector Regression (SVR)[11] applies the same support-vectors based approach from Support Vector Machines to form a flexible tube of minimal radius around the estimated function. The points outside the tube suffer a penalization, but those within the tube will not be penalized. One of the advantages of SVR is that it has excellent generalization capability. Also, its computational complexity does not depend on the dimensionality of the input space. SVR is capable of dealing with non-linear data by simply transforming the data into a different feature space by the so-called kernel trick. Since in the deconvolution problem we assumed it can be explained by a non-linear regression function, we decided to apply a Gaussian/radial kernel to the epsilon-SVR variant: Four parameters were considered during the tuning process:

cost: cost of constraints violation

gamma: the inverse of the radius of influence of samples selected by the model as support vectors.

epsilon: Defines a margin of tolerance where no penalty is given to errors.

Nfeatures: the different number of features according to the rank provided by the Random Forest model.

##### Prediction output

The final model consists of a single sub-model for each cell type, so for instance, in both the coarse and fine grain sub-Challenges we used an SVR model for each of the cell lineages requested, other cell types can be easily added and detected if they are also included in the model. For example, some cell types not included in the challenge were also detected and estimated in our model, like cancer cells or other polymorphonuclear leukocytes.

The input to the prediction resulting model is a vector of RHINO normalized gene expression values (or an expression matrix for multiple samples) and the result is a numeric vector (or matrix) with the estimated cell-type proportion of each cell type in the model. In these regards, it is important to notice that all the cell type models were trained with the same admixture sets, which were bound to contain known purified sample proportions that add up to 1. For instance, one training sample could have 0.2 fibroblast, 0.5 B cells and 0.3 of other cells while another sample could have 0.5 of endothelial cell and 0.5 of cancer cells, which allowed our model to detect different cell type proportions independently one of the other, allowing the possibility of having unseen or unknown cell types in the admixtures without affecting the predicted output and making the predicted proportions comparable across different cell types.

This design should limit the results to have a maximum sum of 1 if there are not unknown cell types present in the mixture but the sum can be less than 1 if otherwise. Of note, no constraints were included in the submitted model but they could be included turning negative prediction to be 0 and rescaling the predictions to 1 if the sum of the predictions is greater than 1, although these alternatives have not been explored thoroughly.

Another important issue to address is that the normalization preprocessing of the training samples and that to perform predictions on new cases is done using the information within each sample independently, which allows the results to be comparable between samples even from different centers, which is of great importance when multi-institution studies are designed or when analyzing isolated samples for the decision-making process in a practical context (i.e. decide treatment in a patient)

These issues must be further explored and specifically addressed by additional research directed at this subject specifically, nevertheless, we believe the rationale of our method is supported by the results obtained using this approach within the current study.

#### Conclusion/Discussion

In conclusion, we believe our method yields a very flexible, yet robust, approach to the tumor deconvolution problem with several desired features that might be of interest for the community:

- It includes bulk RNA-seq experiments from a wide range of sequencing platforms and different research centers that were successfully used to train well-performing models with a reduced set of samples (843 purified samples in total). In these regards, the same approach could be applied with a bigger number of samples possibly improving the model accuracy or it also could be extended to single-cell RNA-Seq experiments, which might be able to detect pure cell types that could be used for training the model.
- It relies on a within-sample normalization procedure that improves the generalization of our method and unlocks the prediction of cell-type proportions from single samples which could be of great importance in future health applications.
- It allows the possibility to include new cell types not studied here.
- It is able to account for unknown cell types in the sample.

- All the model training can be performed on a desktop computer and does not require exceptional computer resources, making it accessible for use by the community and for further research and development.

As future directions, we would like to test our method in real complex tissue samples like tumor samples to evaluate the performance of our method in a real context. As a limitation, we know that immune, cancer and stromal cells suffer from important changes in their phenotype and gene expression patterns in these complex scenarios and that the isolated samples might not represent the true gene expression patterns displayed by the different cell types when they interact with each other.

Also we would like to further validate our proposed normalization procedure RHINO, which we believe could be added to the transcriptomic analysis toolbox.

#### Authors Statement

M Guerrero-Gimenez designed the overall approach and methodology including the normalization procedure. B Lang and M Guerrero-Gimenez curated the datasets and metadata. B Lang downloaded the data, performed sample quality control, trimming and Kallisto aligning. C Catania aided in the overall design and led the machine learning process of the project performing all the model training, hyperparameter tuning, and testing.

### mitten_TDC19 deconvolution method

#### Summary Sentence

We used a score calculated as the sum of the expression of a selected set of markers for each sample.

#### Background/Intro

The key idea of the method was to focus on identifying new markers that are predictive of the relative cell type fractions. This was achieved by first identifying bulk and single-cell datasets of pure cell populations. For single-cell data, we also used non-pure populations and extracted pure populations using previously published methods.[12,13] We then used a Monte Carlo procedure to create random mixtures and identified genes that correlated better with the generated fractions. This took into account the overall distribution of each candidate marker in correlating with the cell fraction.

#### Methods

Fourteen pure cell type RNA-seq datasets were collected from GEO (Table S15). Only control samples, i.e., samples from healthy human donors without treatment, were retained, except for two datasets including samples with breast and colon cancer. Gene expression values were converted to TPM if not in TPM originally. To convert Raw data to TPM, gene length information was obtained from Gencode v31.

We used the above datasets of pure-cell types to create training datasets $T_{jk}$ where $j = 1, ..., m$ are genes and $k = 1, ..., n$ are cell types . We then constructed samples of randomly mixed pure cell type populations, $S_j^a = \sum_k T_{jk} R_k^a$ where $a$ is the in silico sample index with fractions $R_k^a$. The random fractions $R_k^a$ were drawn from a non-uniform distribution enriched for smaller fractions.

Using the $S^a$ ensemble, we calculated the Pearson correlation coefficient between the concentration of cell type $k$ (i.e., $R_k^a$) in the $M_{jk}$ ensemble and the expression of genes. We kept the most correlated genes as $+1$ in a signature matrix , and $0$ otherwise.

In the deconvolution, each dataset $X_{ij}$, where $i$ is a sample and $j$ is a gene, was converted to a linear scale. Each sample $i$ was assigned a score for each cell type $k$ according to $\beta_{ik} = \sum_j X_{ij} M_{jk}$, which was submitted as a prediction.

The same method was used for the two sub-Challenges.

#### Conclusion/Discussion

We focused on the selection of markers and used a basic algorithmic approach for the predictions. The fact that we still ended in the top three performing groups indicates that the selection of markers is probably the most important aspect of this challenge.

#### Authors Statement

SD, TB, and CP designed the algorithms. SD and TB performed calculations. TB submitted models to challenge.

### Biogem deconvolution method

#### Summary Sentence

The method uses robust linear modeling from the R package MASS together with a signature matrix made of harmonized gene expression data normalized by mRNA abundance.

#### Background/Intro

The approach used in this challenge is based on work that was previously published.[14] The feature selection of the approach is done through differential expression and filtering for specific and low-noisy genes. The deconvolution method is robust linear model, which is robust to collinearity and outliers. The signature matrix was normalized by mRNA abundance to take into account the different mRNA yields of the various immune cell types.

The main variation in respect to the published paper[14] is the utilization of a large harmonized dataset for the generation of the signature matrix. The benefit of this approach over other popular approaches is that it allows to achieve top level results without the utilization of more complex algorithms based on machine and deep learning.

#### Methods

##### Data collection and preprocessing

The publicly available data used to generate the signature matrix were taken from the following datasets: PRJEB14751, PRJEB36933, PRJNA218851, PRJNA272556, PRJNA327180, PRJNA338944, PRJNA352224, PRJNA418779, PRJNA430418, PRJNA449980, PRJNA471906, PRJNA483877, PRJNA484735, PRJNA489270, PRJNA490870, PRJNA495625, PRJNA540256, PRJNA559359, PRJNA562113, PRJNA598222 (Table S16).

The following samples were selected independently of the condition or tissue they were derived from and for some datasets only a subset of samples were downloaded and processed for this challenge:

| Dataset | Cell type | Condition | Platform | Submissions |
|---|---|---|---|---|
| PRJEB14751 | Endothelial cells | HUVEC cell line | HiSeq 2500 | 1, 2, 3 |
| PRJEB36933 | Tregs | healthy donors | HiSeq 4000 | 2, 3 |
| PRJNA218851 | CRC | primary tumor | HiSeq 2000 | 1, 2, 3 |
| PRJNA272556 | CRC | cell lines | HiSeq 1000 | 1, 2, 3 |
| PRJNA327180 | Fibroblasts | cell lines | HiSeq 2000 | 1, 2, 3 |
| PRJNA338944 | Fibroblasts | human cancer | HiSeq 2500 | 1, 2, 3 |
| PRJNA352224 | mDCs | healthy stimulated and non | HiSeq 2000 | 2, 3 |
| PRJNA418779 | Various immune | healthy | HiSeq 2000 | 1, 2, 3 |
| PRJNA430418 | Endothelial cells | stimulated and non | HiSeq 2500 | 1, 2, 3 |
| PRJNA449980 | Macrophages | healthy simulated and non | HiSeq 2500 | 3 |
| PRJNA471906 | Endothelial cells | healthy | HiSeq 2500 | 1, 2, 3 |
| PRJNA483877 | Monocytes | breast cancer | HiSeq 2500 | 1, 2 |
| PRJNA484735 | Various immune | healthy stimulated and non | NovaSeq 6000 | 1, 2, 3 |
| PRJNA489270 | BRCA | cell lines | HiSeq 2000 | 1, 2, 3 |
| PRJNA490870 | CRC | primary tumor | HiSeq 4000 | 1, 2, 3 |
| PRJNA495625 | BRCA | cell lines | HiSeq 2000 | 1, 2, 3 |
| PRJNA540256 | Macrophages | from endometrium | HiSeq 4000 | 1, 2 |
| PRJNA559359 | Macrophages | healthy | HiSeq 2500 | 3 |
| PRJNA562113 | CRC | primary and metastatic tumor | HiSeq 2500 | 1, 2, 3 |
| PRJNA598222 | BRCA | primary tumor | HiSeq 2000 | 1, 2, 3 |

The fastq files were downloaded from the ENA archive and kallisto (version 0.46.2)[6,14] was used to obtain gene expression data. The pseudo-alignment was done against the GENCODE transcriptome (release 33, genome assembly GRCh38).

#### Feature selection

The first feature selection step consists in performing differential expression analysis between each cell type and remaining ones. This was done using the voom and limma methods.[8]

The second feature selection step consists in filtering out genes that are not beneficial for deconvolution. These are:

1. highly expressed genes -- any gene that has a TPM value > 3000 in at least one sample;
2. low expressed genes -- any gene with a summed up expression value < 5 across all samples;

3. specificity -- any gene *g* from the differential analysis between cell type *c* and the remaining cell types that has an effect size < 0.1 between its median $\log_2$ values in *c* and the cell type *c*' with the second highest expression (after *c*) of *g*.

The third feature selection step consists of a hard threshold filter to avoid the over-representation of cell types with many remaining genes. Hence, no more than 70 genes and 90 genes were kept for each cell type for the Coarse and Fine sub-Challenges, respectively.

##### Signature matrix

For the signature matrix, for each cell type the median value of counts per million (CPM) was generated. Next, the values were scaled by a factor that accounts for mRNA abundance.[14] The previously published[14] scaling factors were only for cell types from PBMCs. Hence, the scaling factors for tumor cells, endothelial cells and fibroblasts were conventionally set to 1.

##### Prediction method

The prediction method used for deconvolution is the Robust Linear Model, which was implemented using the rlm function from the R package MASS. The method is robust to noise and collinearity and its performance has been shown to be comparable to support vector regression as used in CIBERSORT.[14,15]

The method expects CPM as input and the code includes a step that checks that the sum of every sample of the input data is 10^6.

##### Prediction output

The values generated by the robust linear model are interpretable as fractions.

The negative values are dealt with in the following two steps: 1) any value lower than -2 is set as -2; 2) scale values so that the minimum value is 0.

The output values can be used for both comparison between and within samples.

To allow comparison between samples, the approach does not implement any constraints. In this way, if the sample has an amount of unknown content that is not predicted by the method, there is no inflation of the fractions due to a method constraint.

To allow comparison within samples, the approach implements normalization of mRNA abundance to the signature matrix.[14] Normalization for mRNA abundance takes into consideration the higher mRNA content of certain cell types (such as monocytes) and thereby avoids inflation of their predicted fractions.

Therefore, this method allows the deconvolution of absolute abundance of immune cell types present in the mixture sample.

#### Variations in 2nd and 3rd submission

The variation included in the 2nd and 3rd submission do not affect the method, but only the implementation of batch effect correction and the inclusion and exclusion of various datasets.

For the 2nd and 3rd submission, the outlier samples to exclude were evaluated visually through a PCA analysis and were excluded from subsequent analyses. The data were also corrected for batch effects using Combat.

Fig. S3 shows that after batch effect correction of some immune cell datasets, the samples do not cluster according to dataset anymore.

Regarding specific datasets that allowed us to achieve better performance, we noticed that we could improve the scores obtained on T regulatory cells only after adding the dataset PRJEB36933. Moreover, to improve the results obtained for macrophages, we excluded the samples from the datasets PRJNA483877 and PRJNA540256, and we added samples from the datasets PRJNA449980 and PRJNA559359.

We used the following number of samples per cell type in the coarse-grained sub-Challenge:

| Cell Type | Submission 1 | Submission 2 | Submission 3 |
|---|---|---|---|
| B.cells | 38 | 38 | 38 |
| BRCA | 10 | 10 | 10 |
| CD4.T.cells | 88 | 100 | 98 |
| CD8.T.cells | 51 | 51 | 51 |
| CRC | 20 | 20 | 20 |
| endothelial.cells | 19 | 19 | 19 |
| fibroblasts | 10 | 10 | 10 |
| monocytic.lineage | 58 | 61 | 53 |
| neutrophils | 4 | 4 | 4 |
| NK.cells | 15 | 15 | 15 |
| SUM | 313 | 328 | 318 |

We used the following number of samples per cell type in the fine-grained sub-Challenge:

| Cell Type | Submission 1 | Submission 2 | Submission 3 |
|---|---|---|---|
| BRCA | 10 | 10 | 10 |
| CRC | 20 | 20 | 20 |
| endothelial.cells | 19 | 19 | 19 |
| fibroblasts | 10 | 10 | 10 |
| macrophages | 22 | 22 | 20 |
| memory.B.cells | 19 | 19 | 19 |
| memory.CD4.T.cells | 40 | 40 | 40 |
| memory.CD8.T.cells | 27 | 27 | 27 |
| monocytes | 29 | 29 | 18 |
| myeloid.dendritic.cells | 7 | 10 | 15 |
| naive.B.cells | 12 | 12 | 12 |
| naive.CD4.T.cells | 4 | 4 | 4 |
| naive.CD8.T.cells | 12 | 12 | 12 |
| neutrophils | 4 | 4 | 4 |
| NK.cells | 15 | 15 | 15 |
| regulatory.T.cells | 23 | 33 | 33 |
| SUM | 273 | 286 | 278 |

Through these variations, a substantial improvement of the approach was achieved for the fine challenge, although not much for the coarse challenge.

##### Variations between sub-Challenge 1 (coarse-grained sub-Challenge) and sub-Challenge 2 (fine-grained sub-Challenge)

The data processing and methodology used for the fine-grained sub-Challenge are identical to the ones for the coarse-grained sub-Challenge.

There are only these two consideration to make:

- regarding the normalization for mRNA abundance, for macrophages we used the same scaling factor calculated for classical monocytes as they belong to the same lineage.
- for the hard threshold of the number of genes to include for cell type, we kept no more than 70 genes for the coarse-grained sub-Challenge and no more than 90 genes for the fine-grained sub-Challenge.

#### Conclusion/Discussion

In conclusion, we believe we reached a good deconvolution performance using robust linear modeling on a signature matrix generated from a large harmonized dataset. We believe that more datasets are necessary to assess the variability across certain cell types and to establish which isolation strategies or biological conditions increase such variability. This is especially the case if one is trying to perform deconvolution on subtypes of the same lineage, such as memory T cell subtypes.

#### Authors Statement

GM collected, processed the samples, and developed the methodology. FPC contributed to data collection and data analysis. MC contributed in the interpretation of the results.

#### IZI deconvolution method

The method aims to utilize a probabilistic description of the cell-type specific transcription composition. We gathered thousands of replicates for gene expression measurements per cell type from 75,720 different samples in hundreds of publicly available GEO datasets. The replicates include single cell RNA-seq, bulk RNA-seq and bulk expression microarray data. To harmonize the datasets, we selected genes and samples such that the amount of recovered samples and genes that are shared between all samples are maximized. This resulted in a total of 21,992 harmonized RNA-expression vectors for the coarse-grained sub-Challenge and 13,317 for the fine-grained sub-Challenge. We then applied a pipeline of transformations and dimensional reduction that are both reversible and differentiable to the expression data. The goal was to find transcriptome representations such that the distribution per cell type can be described through normal distributions in a latent space where the Wasserstein distance between pairs of cell types is maximized. We assumed that the result of the RNA-seq experiment and counting process is a random process that is influenced by the composition of transcripts in the sequenced sample. Specifically, we assume that finding a read for a given sample is a Bernoulli experiment with a fixed probability for each sample. Bayes theorem gives rise to a probabilistic description of the transcriptome composition based on the raw observed counts through the Dirichlet distribution. The decomposition space can be transformed with an isometric log-ratio transformation to achieve approximate normal distributions. Through scaling such that the average covariance per cell type is the unit matrix and applying principal component analysis on the cell-type means, we found a dimensional reduction that approximately maximizes the Wasserstein distance between the cell-type specific distributions. A justification of this procedure has been previously published.[16] The resulting distributions in dimensionally reduced space serve as characterization of the respective cell types and define a distribution of transcriptome compositions in expression space. We used these characterizations to define a Bayesian model that mixes the cell-type-specific transcriptomes with Dirichlet distributed weights to describe the expression profile of the mixed samples that ought to be deconvolved. Through the application of automatic variational differential inference (ADVI)[17] in the pymc3 package[18] we performed Bayesian inference without exceeding the computation resource constraints and use the posterior mean of the mixture weights as an output for the deconvolution. The algorithm aims to make very little assumptions about the data and could potentially achieve absolute cell mass quantifications of individual samples without knowing the context of the cohort. The main limitations are the approximation of the posterior through ADVI and inaccuracies of the cell type characterization through mislabeling or lack of transcriptome samples, e.g., we used only 42 memory CD8 T-cells.

### Supplemental Figures

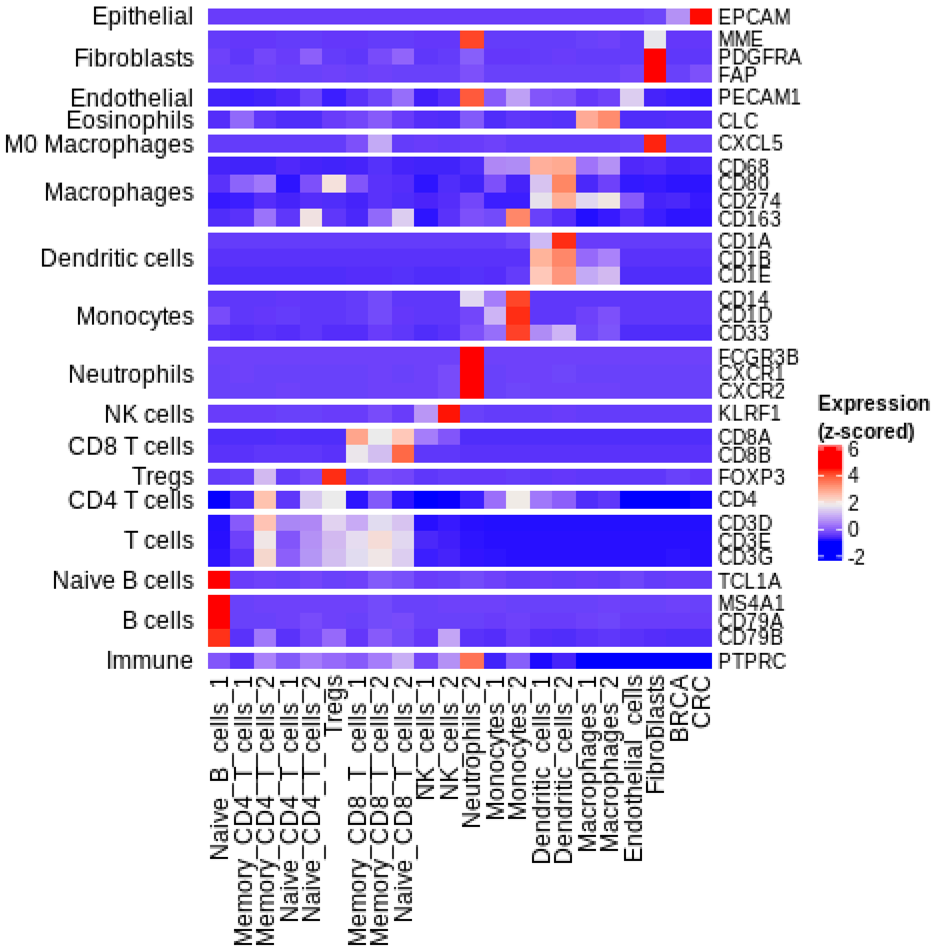


**Fig. S1:** Marker gene expression within purified populations. Expression of markers (right axis) grouped by corresponding cell population (left axis) within each purified sample (columns).

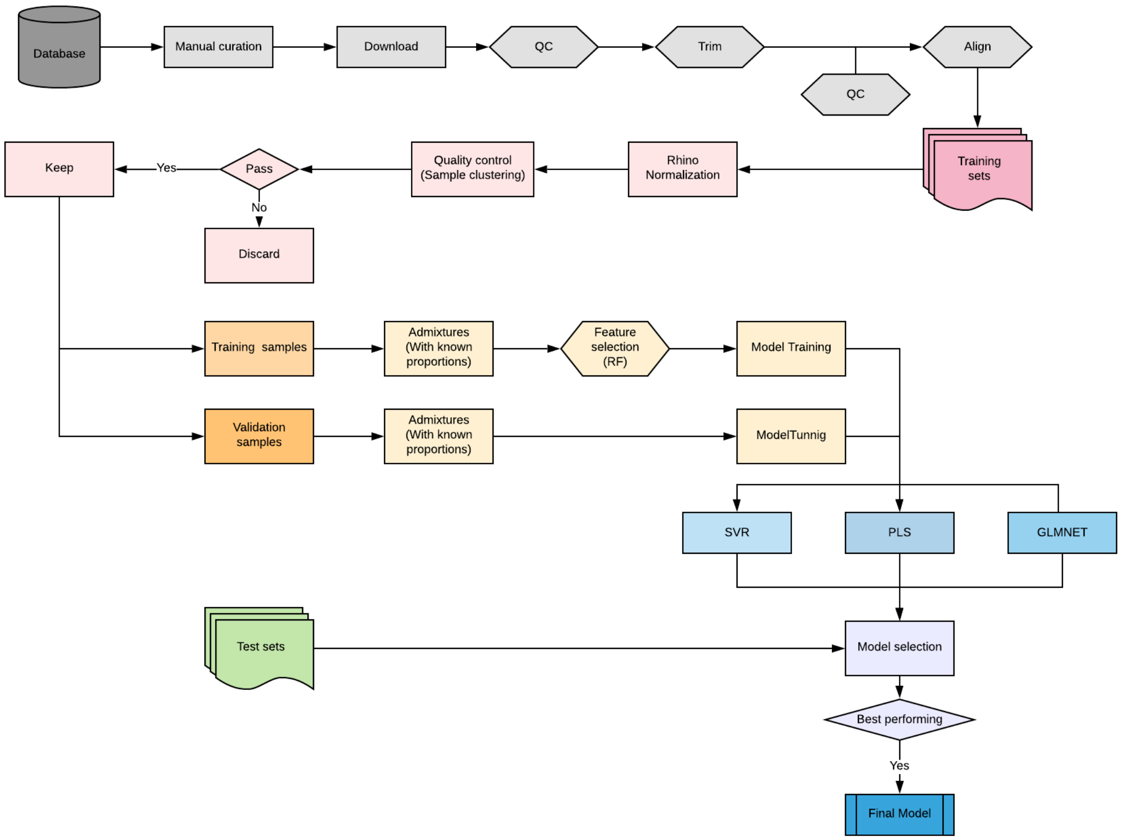


**Fig. S2:** Workflow of DA_505 deconvolution method.

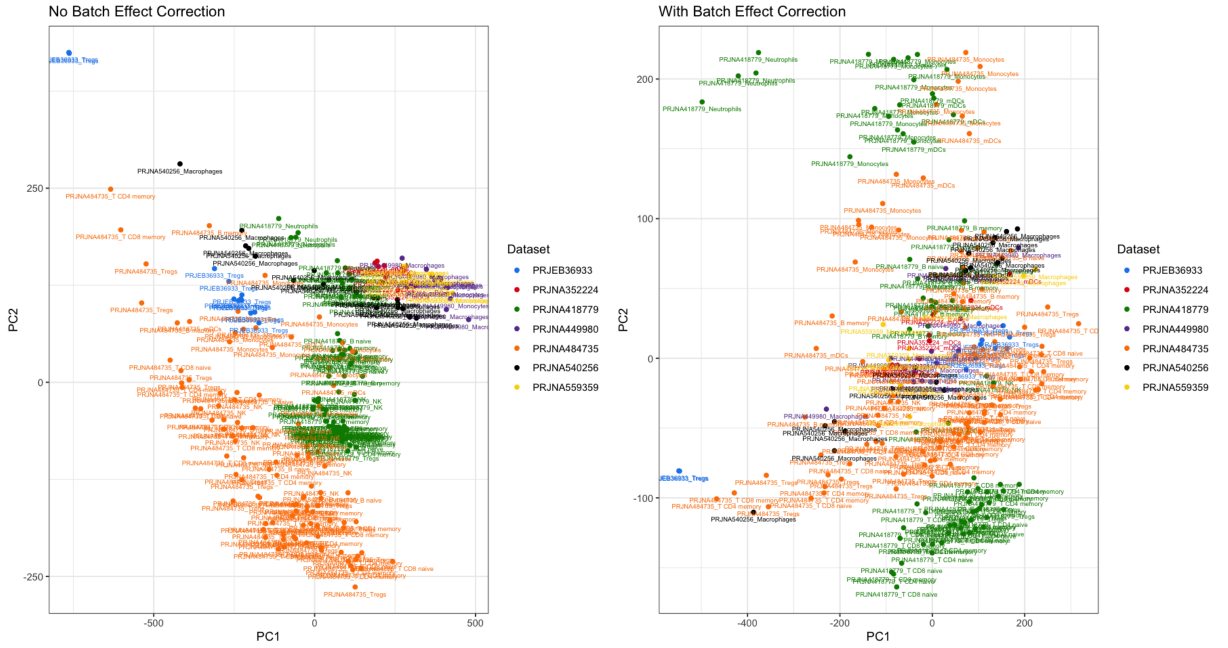


**Fig. S3:** Batch correction of data used to train Biogem deconvolution method.

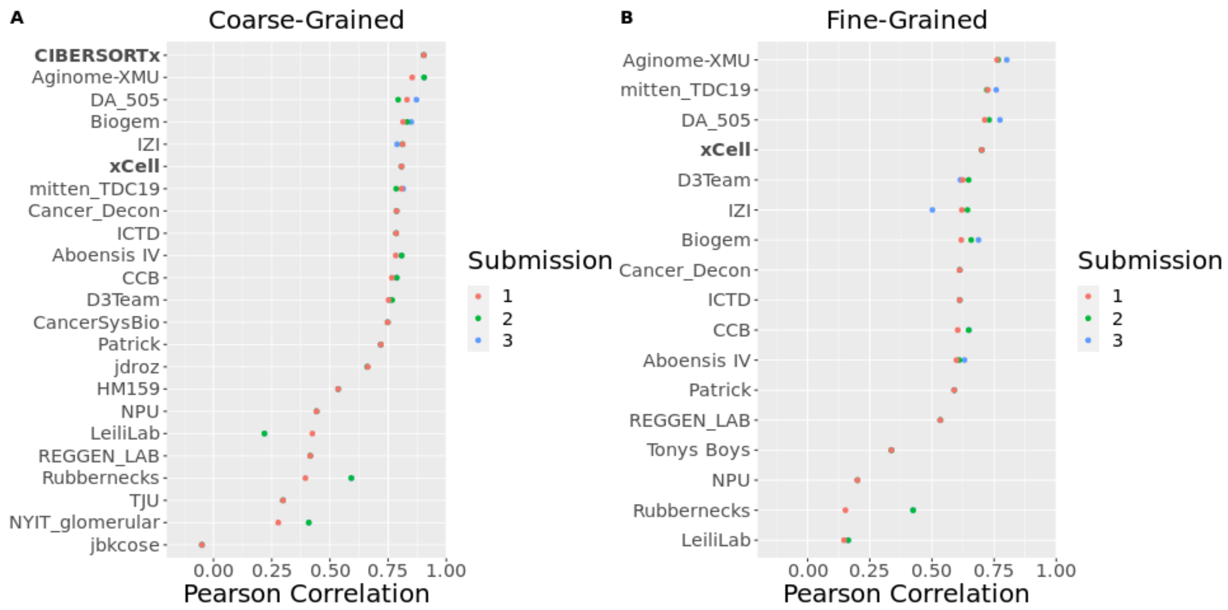


**Fig. S4:** Aggregate primary, Pearson-based score of participant methods over submissions and of comparator methods. (A, B) Aggregate Pearson-based score of methods in (A) coarse- and (B) fine-grained sub-Challenges over bootstraps (*n*=1,000; Methods). Comparator methods (bold) are shown only if their published reference signatures include all cell types in each respective sub-Challenge: CIBERSORTx (coarse-grained only) and xCell. Methods ordered by Pearson correlation of first submission in respective sub-Challenge.

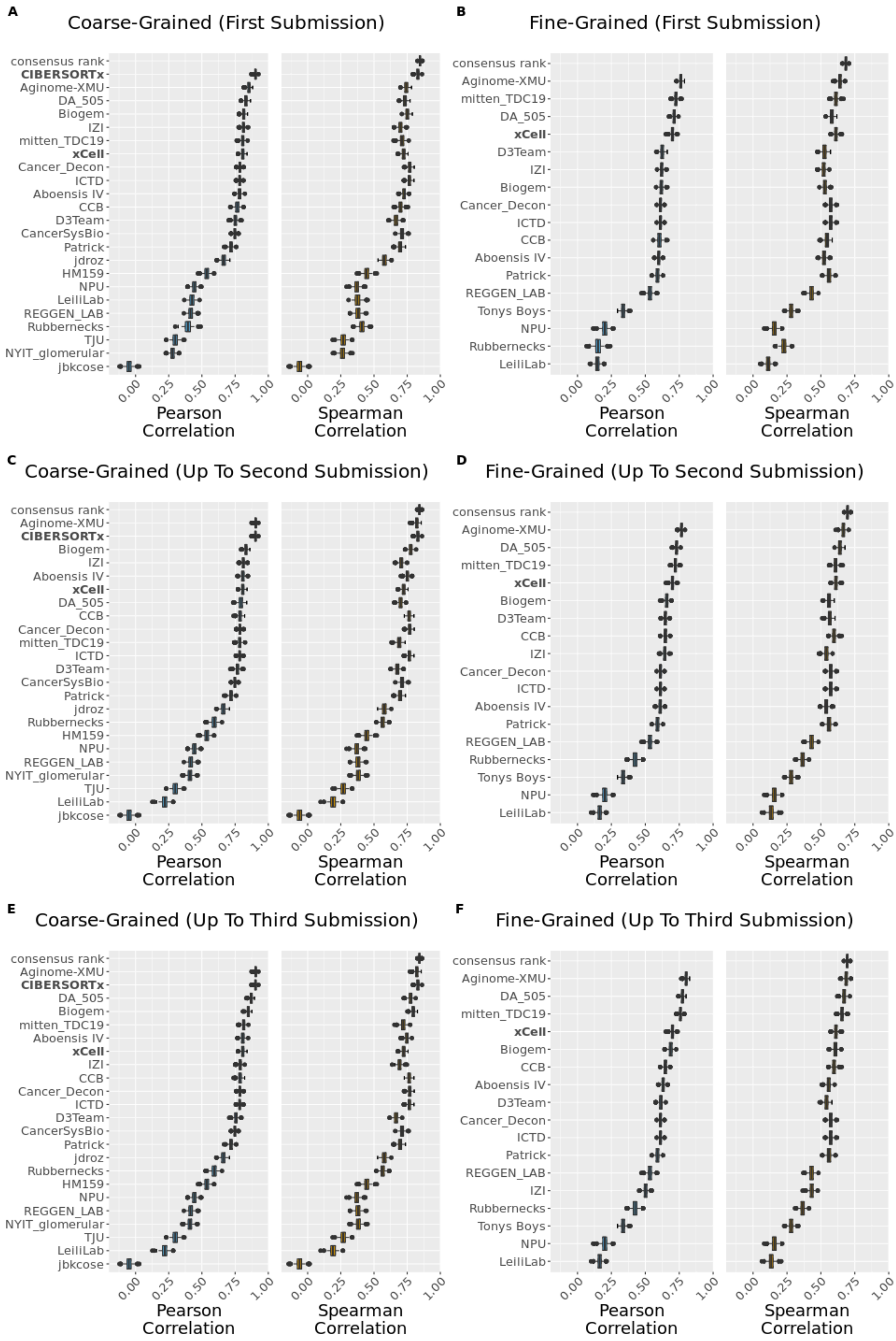
A
Coarse-Grained (First Submission)
consensus rank
CIBERSORTx
Aginome-XMU
DA_505
Biogem
IZI
mitten_TDC19
xCell
Cancer_Decon
ICTD
Aboensis IV
CCB
D3Team
CancerSysBio
Patrick
jdroz
HM159
NPU
LeiliLab
REGGEN_LAB
Rubbernecks
TJU
NYIT_glomerular
jbkcose
Pearson Correlation
Spearman Correlation
0.00 0.25 0.50 0.75 1.00
B
Fine-Grained (First Submission)
consensus rank
Aginome-XMU
mitten_TDC19
DA_505
xCell
D3Team
IZI
Biogem
Cancer_Decon
ICTD
CCB
Aboensis IV
Patrick
REGGEN_LAB
Tonys Boys
NPU
Rubbernecks
LeiliLab
C
Coarse-Grained (Up To Second Submission)
consensus rank
Aginome-XMU
CIBERSORTx
Biogem
IZI
Aboensis IV
xCell
DA_505
CCB
Cancer_Decon
mitten_TDC19
ICTD
D3Team
CancerSysBio
Patrick
jdroz
Rubbernecks
HM159
NPU
REGGEN_LAB
NYIT_glomerular
TJU
LeiliLab
jbkcose
D
Fine-Grained (Up To Second Submission)
consensus rank
Aginome-XMU
DA_505
mitten_TDC19
xCell
Biogem
D3Team
CCB
IZI
Cancer_Decon
ICTD
Aboensis IV
Patrick
REGGEN_LAB
Rubbernecks
Tonys Boys
NPU
LeiliLab
E
Coarse-Grained (Up To Third Submission)
consensus rank
Aginome-XMU
CIBERSORTx
DA_505
Biogem
mitten_TDC19
Aboensis IV
xCell
IZI
CCB
Cancer_Decon
ICTD
D3Team
CancerSysBio
Patrick
jdroz
Rubbernecks
HM159
NPU
REGGEN_LAB
NYIT_glomerular
TJU
LeiliLab
jbkcose
F
Fine-Grained (Up To Third Submission)
consensus rank
Aginome-XMU
DA_505
mitten_TDC19
xCell
Biogem
CCB
Aboensis IV
D3Team
Cancer_Decon
ICTD
Patrick
REGGEN_LAB
IZI
Rubbernecks
Tonys Boys
NPU
LeiliLab

**Fig. S5**: Aggregate method performance over rounds. Aggregate score (primary metric: Pearson correlation; secondary metric: Spearman correlation) of participant and comparator methods in (A, C, E) coarse- and (B, D, F) fine-grained sub-Challenges over bootstraps (*n*=1,000; Methods). Scores reported from the (A, B) first submission, (C, D) second submission (or latest submission up to the second, if less than two submissions), or (E, F) third submission (or latest submission up to the third, if less than three submissions). Comparator methods (bold) are shown only if their published reference signatures include all cell types in each respective sub-Challenge: CIBERSORTx (coarse-grained only) and xCell. Boxplots display median (center line), 25th and 75th percentiles (hinges), and 1.5x interquartile range (whiskers). Methods ordered by median Pearson correlation in respective sub-Challenge. Source data are provided as a Source Data file.

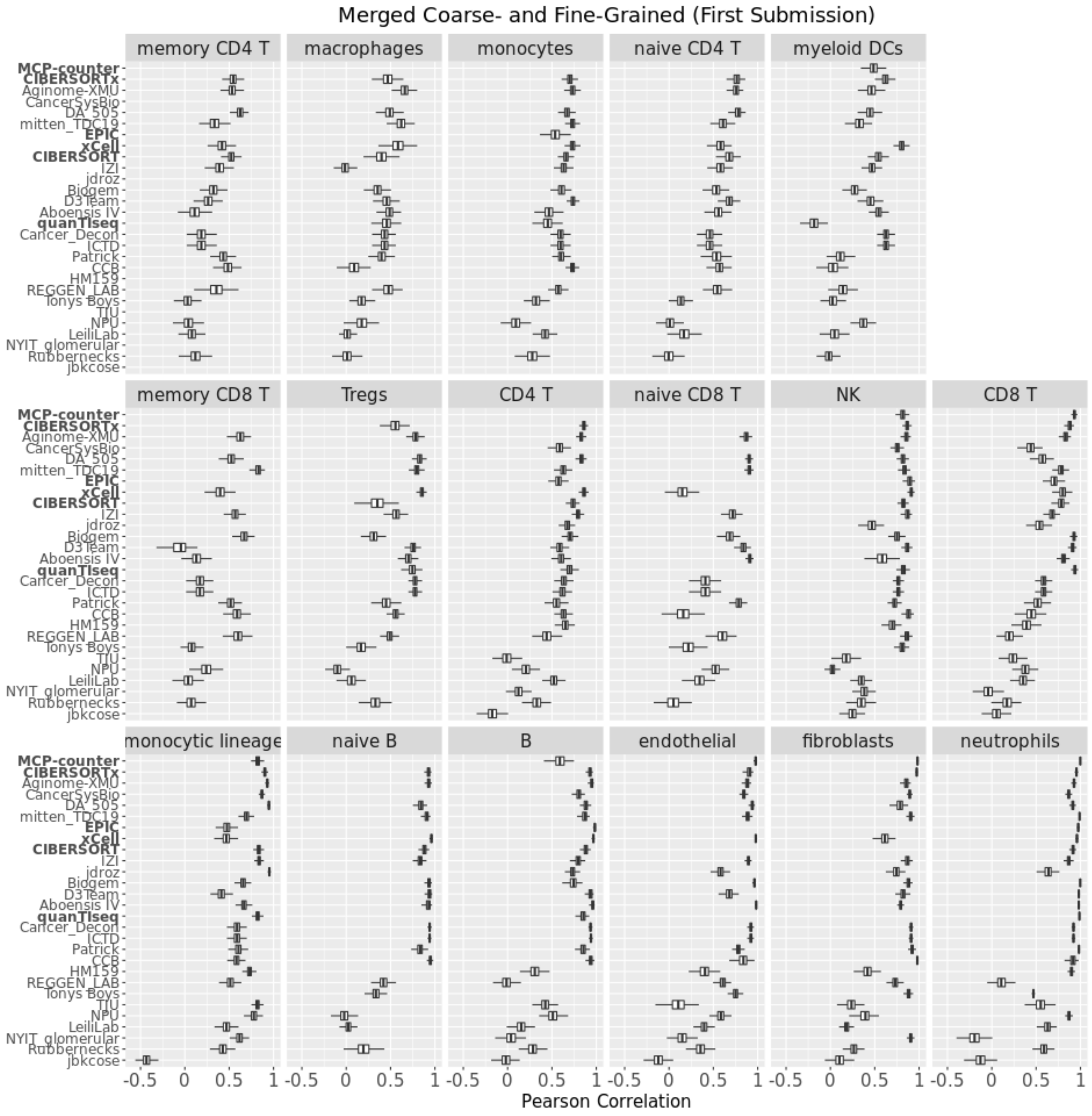


**Fig. S6**: Distribution of per-cell type method performance from first submission merged across coarse- and fine-grained sub-Challenges. Performance (Pearson correlation; x axis) of comparator baseline methods (bold) and participant methods (y axis) for each cell type (facet label). Distribution of Pearson correlations over bootstraps (*n*=1,000; Methods), computed as average over validation datasets and subsequently over coarse- and fine-grained sub-Challenges for cell types occurring in both. Blank row indicates cell type not reported by the corresponding method. Boxplots display median (center line), 25th and 75th percentiles (hinges), and 1.5x interquartile range (whiskers). Methods ordered according to their mean Pearson correlation across cell types (mean column in Fig. 3), and cell types ordered according to their max Pearson correlation across methods (max row in Fig. 3). Source data are provided as a Source Data file.

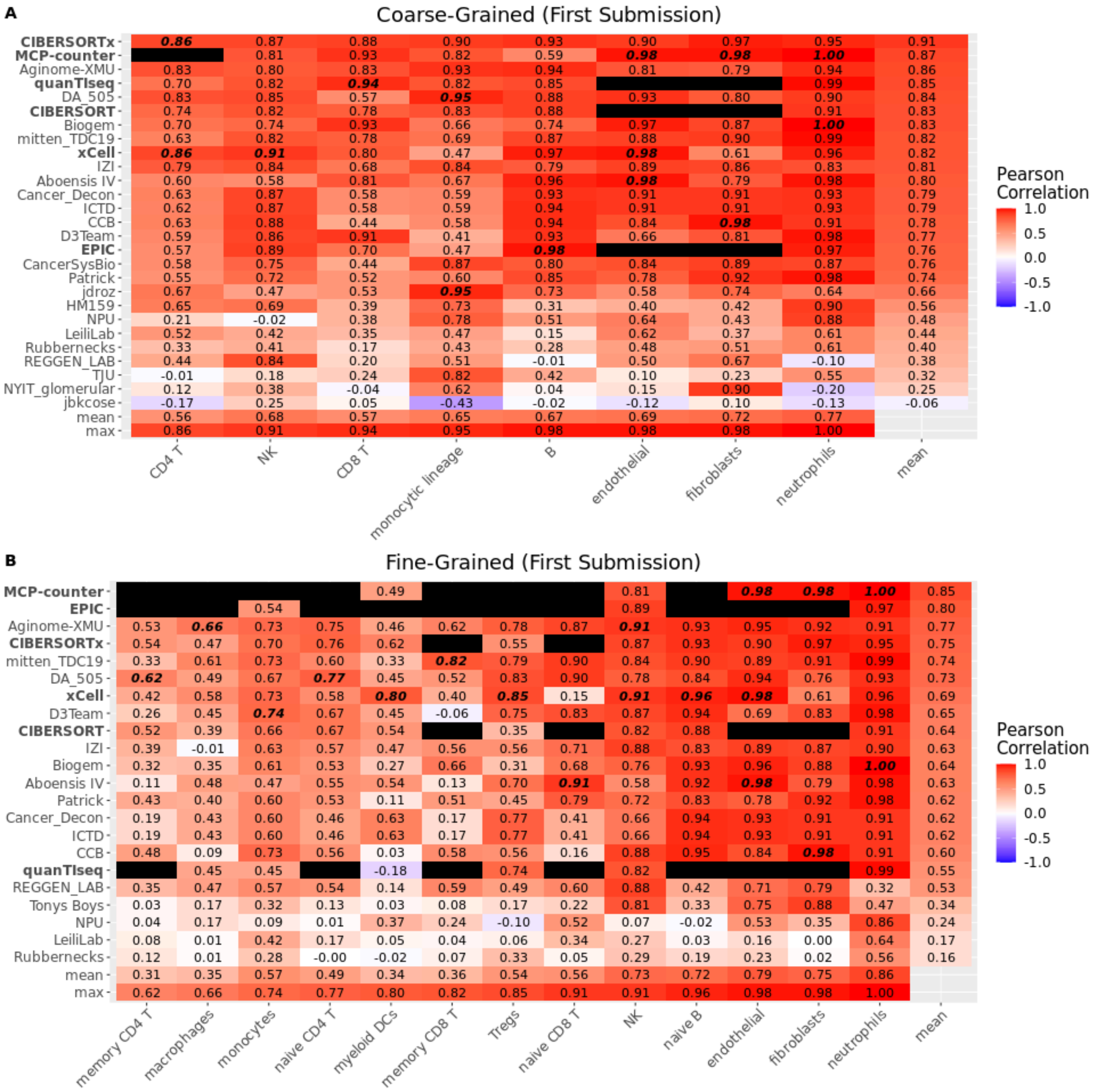


**Fig. S7**: Per-cell type method performance from first submission stratified by sub-Challenge. Pearson correlation of method (left axis) prediction versus known proportion from admixture for each cell type (bottom axis) in (A) coarse- or (B) fine-grained sub-Challenge. Pearson correlation first averaged over validation dataset and then (*n*=1,000) over bootstraps (Methods). Black entry indicates cell type not predicted by corresponding method. Bottom two rows are the mean and maximum correlation, respectively, for corresponding cell type across methods. Rightmost column is mean correlation for corresponding method across predicted cell types. Highest correlation for each cell type highlighted in bold italics. Comparator methods in bold. Methods ordered according to their mean Pearson correlation across cell types (rightmost column), and cell types ordered according to their maximum Pearson correlation across methods (bottom row). Source data are provided as a Source Data file.

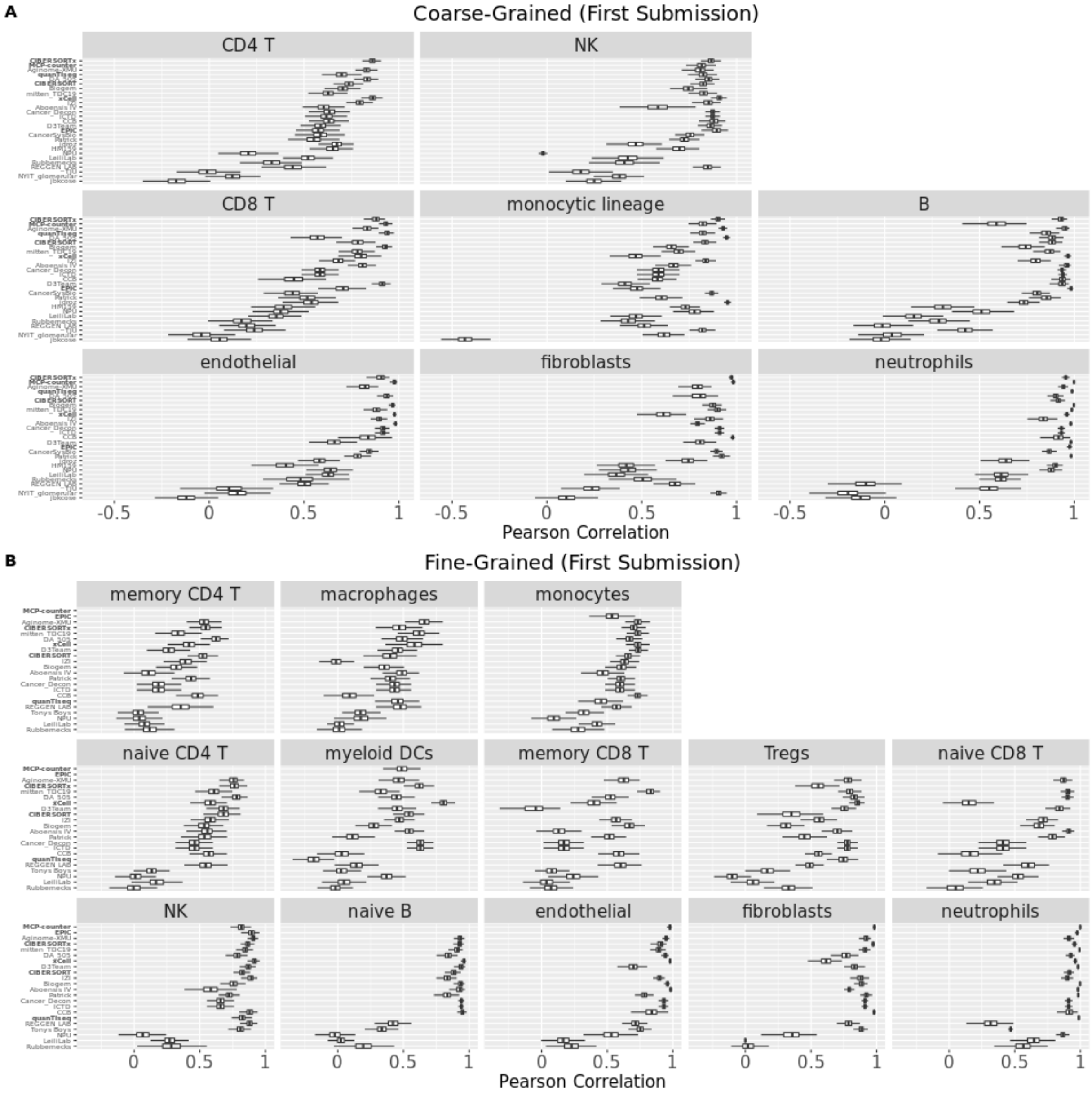


**Fig. S8**: Distribution of per-cell type method performance from first submission stratified by sub-Challenge. Performance (Pearson correlation; x axis) in (A) coarse- and (B) fine-grained sub-Challenges of comparator baseline methods (bold) and participant methods (y axis) for each cell type (facet label). Distribution of Pearson correlations over bootstraps (*n*=1,000; Methods). Blank row indicates cell type not reported by the corresponding method. Boxplots display median (center line), 25th and 75th percentiles (hinges), and 1.5x interquartile range (whiskers). Methods ordered according to their mean Pearson correlation across cell types (corresponding rightmost column of Fig. S7), and cell types ordered according to their maximum Pearson correlation across methods (corresponding bottom row of Fig. S7). Source data are provided as a Source Data file.

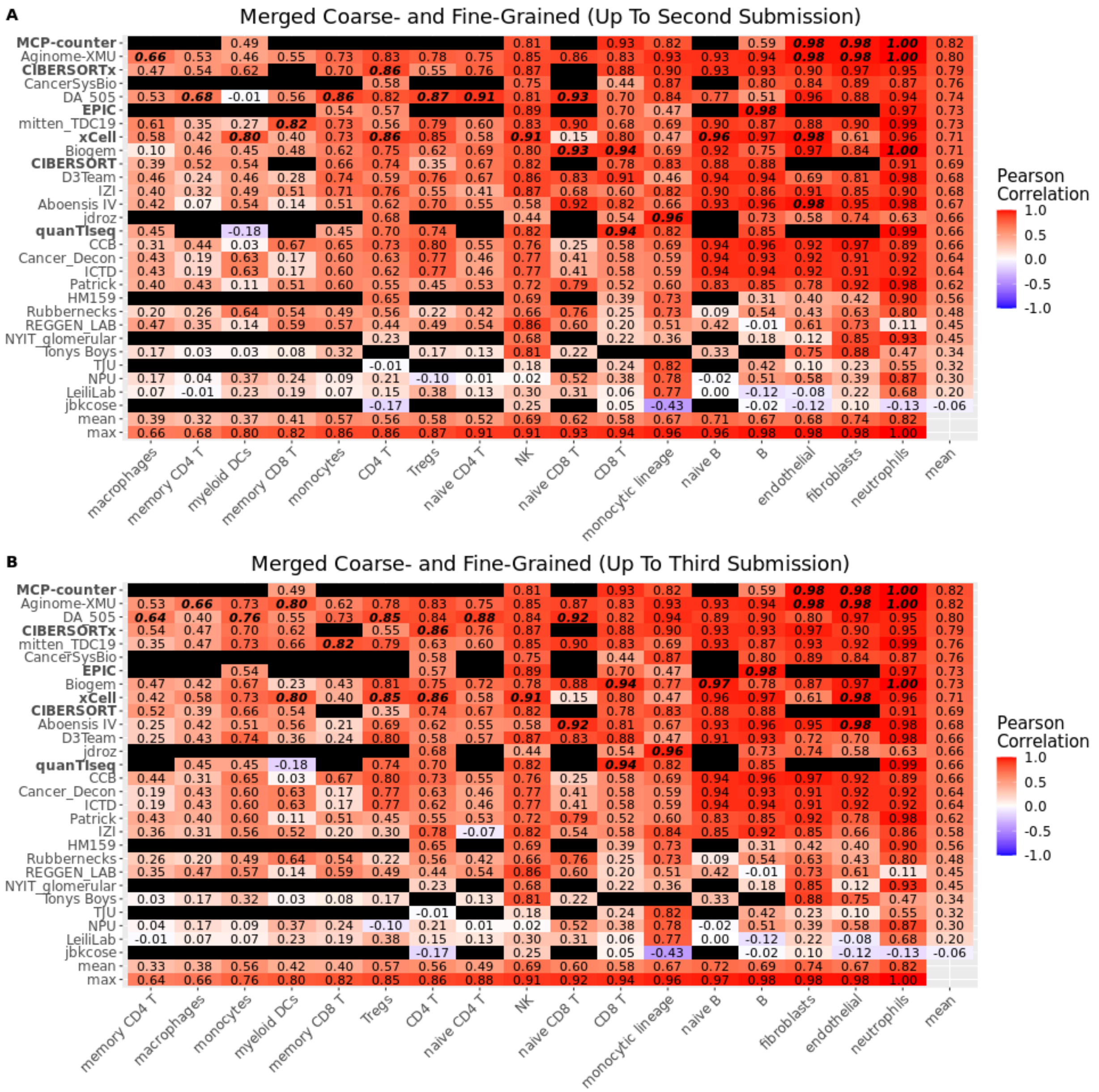

**Fig. S9**: Per-cell type method performance from second and third submissions merged across coarse- and fine-grained sub-Challenges. Pearson correlation of method (left axis) prediction versus known proportion from admixture for each cell type (bottom axis) for (A) second or (B) third submission. Pearson correlation first averaged over validation dataset and then ($n$=1,000) over bootstraps (Methods). Results from latest submission up to (A) second or (B) third submission for those methods with fewer than two or three submissions, respectively.  Black entry indicates cell type not predicted by corresponding method. Bottom two rows are the mean and maximum correlation, respectively, for corresponding cell type across methods.  Rightmost column is mean correlation for corresponding method across predicted cell types. Highest correlation for each cell type highlighted in bold italics. Comparator methods in bold. Methods ordered according to their mean Pearson correlation across cell types (rightmost column), and cell types ordered according to their maximum Pearson correlation across methods (bottom row). Source data are provided as a Source Data file.

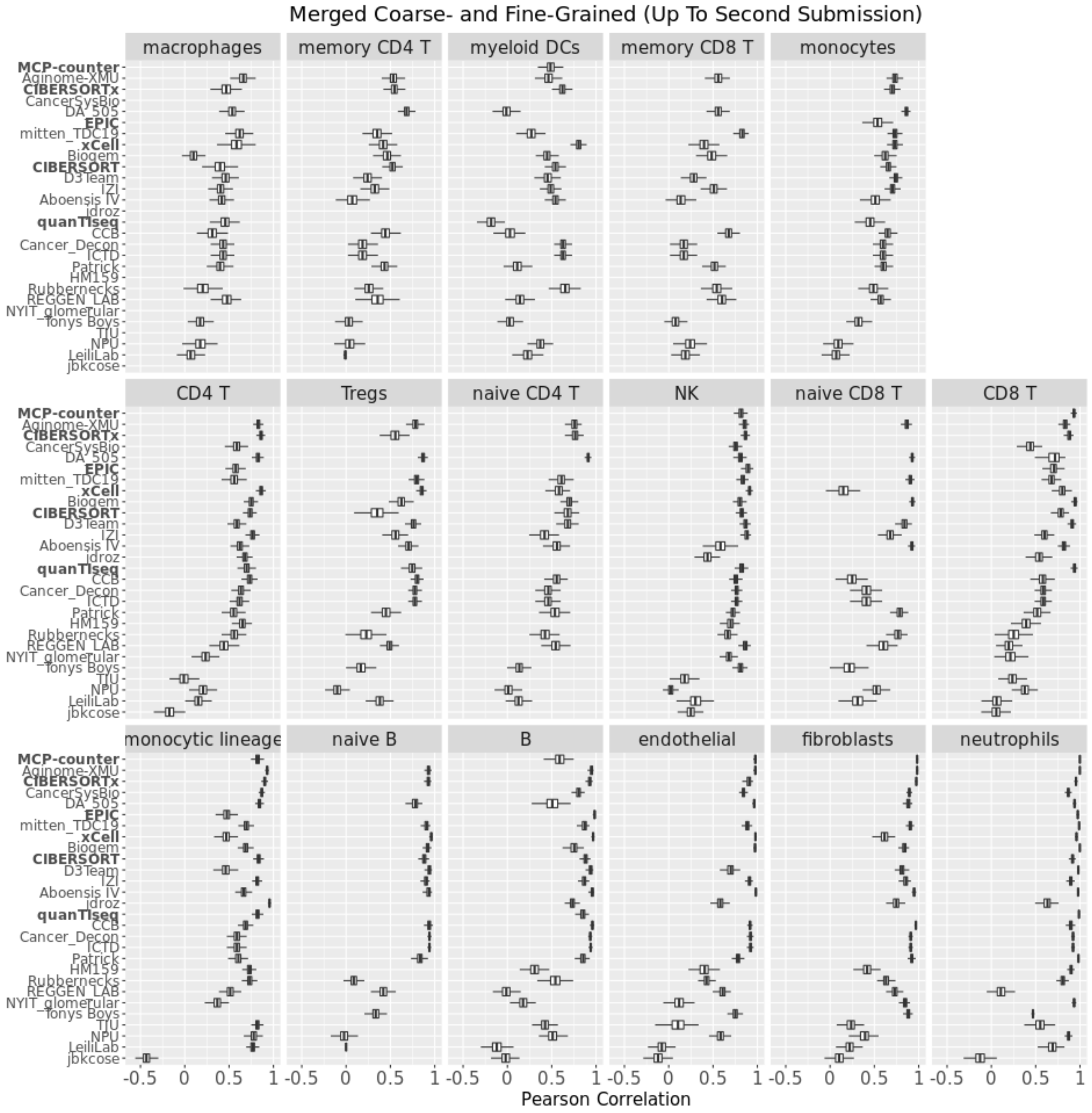


**Fig. S10**: Distribution of per-cell type method performance from second submission merged across coarse- and fine-grained sub-Challenges. Performance (Pearson correlation; x axis) of comparator baseline methods (bold) and participant methods (y axis) for each cell type (facet label). Distribution of Pearson correlations over bootstraps (*n*=1,000; Methods), computed as average over validation datasets and subsequently over coarse- and fine-grained sub-Challenges for cell types occurring in both. Results from latest submission up to the second, if less than two submissions for corresponding method. Blank row indicates cell type not reported by the corresponding method. Boxplots display median (center line), 25th and 75th percentiles (hinges), and 1.5x interquartile range (whiskers). Methods ordered according to their mean Pearson correlation across cell types (rightmost column of Fig. S9A), and cell types ordered according to their maximum Pearson correlation across methods (bottom row of Fig. S9A). Source data are provided as a Source Data file.

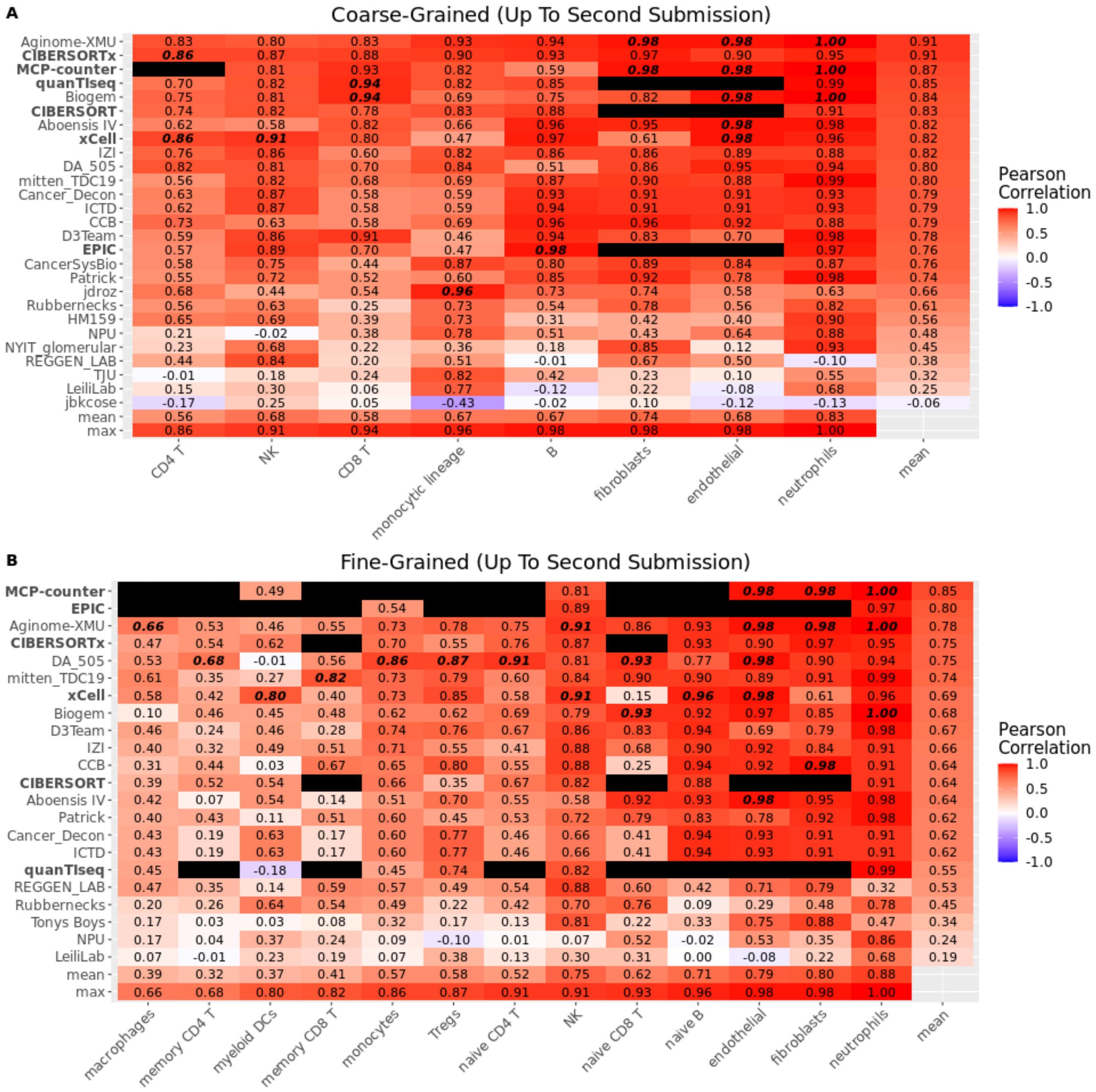


**Fig. S11**: Per-cell type method performance from second submission stratified by sub-Challenge. Pearson correlation of method (left axis) prediction versus known proportion from admixture for each cell type (bottom axis) in (A) coarse- or (B) fine-grained sub-Challenge. Pearson correlation first averaged over validation dataset and then ($n$=1,000) over bootstraps (Methods). Results from latest submission up to second for those methods with fewer than two submissions. Black entry indicates cell type not predicted by corresponding method. Bottom two rows are the mean and maximum correlation, respectively, for corresponding cell type across methods. Rightmost column is mean correlation for corresponding method across predicted cell types. Highest correlation for each cell type highlighted in bold italics. Comparator methods in bold. Methods ordered according to their mean Pearson correlation across cell types (rightmost column), and cell types ordered according to their maximum Pearson correlation across methods (bottom row). Source data are provided as a Source Data file.

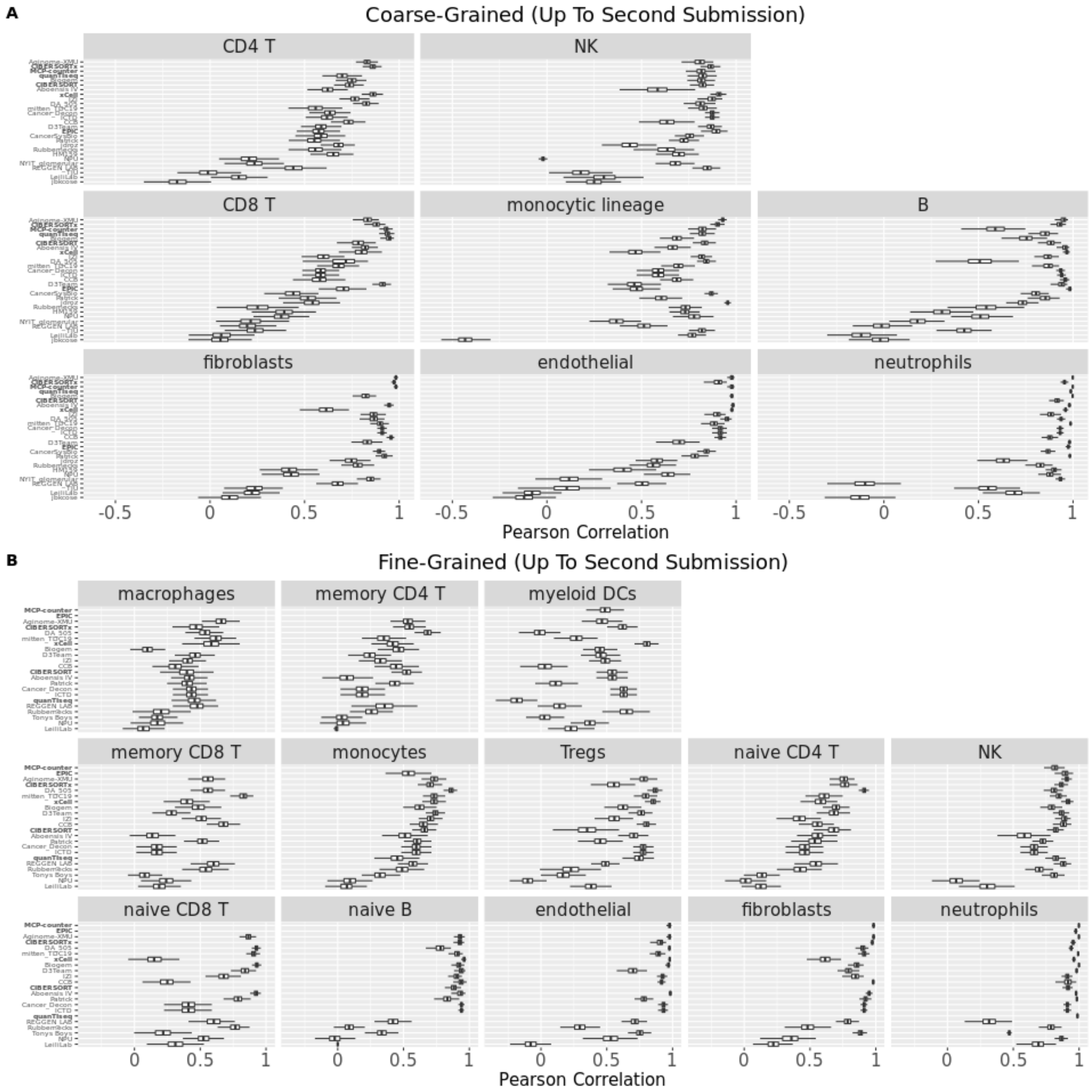


**Fig. S12**: Distribution of per-cell type method performance from second submission stratified by sub-Challenge. Performance (Pearson correlation; x axis) in (A) coarse- and (B) fine-grained sub-Challenges of comparator baseline methods (bold) and participant methods (y axis) for each cell type (facet label). Distribution of Pearson correlations over bootstraps (*n*=1,000; Methods). Results from latest submission up to the second, if less than two submissions for corresponding method. Blank row indicates cell type not reported by the corresponding method. Boxplots display median (center line), 25th and 75th percentiles (hinges), and 1.5x interquartile range (whiskers). Methods ordered according to their mean Pearson correlation across cell types (corresponding rightmost column of Fig. S11), and cell types ordered according to their maximum Pearson correlation across methods (corresponding bottom row of Fig. S11). Source data are provided as a Source Data file.

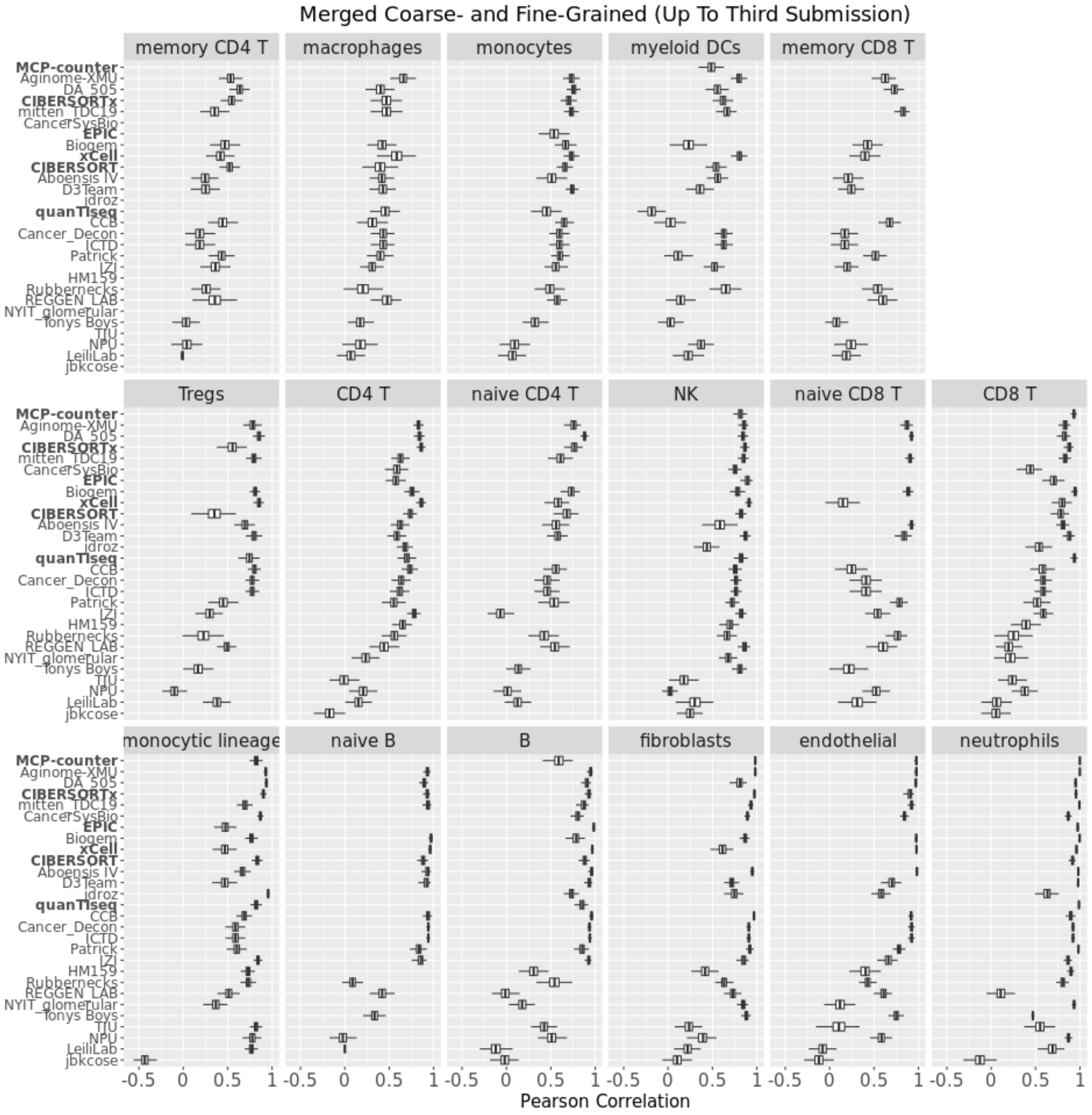


**Fig. S13**: Distribution of per-cell type method performance from third submission merged across coarse- and fine-grained sub-Challenges. Performance (Pearson correlation; x axis) of comparator baseline methods (bold) and participant methods (y axis) for each cell type (facet label). Distribution of Pearson correlations over bootstraps (*n*=1,000; Methods), computed as average over validation datasets and subsequently over coarse- and fine-grained sub-Challenges for cell types occurring in both. Results from latest submission up to the third, if less than three submissions for corresponding method. Blank row indicates cell type not reported by the corresponding method. Boxplots display median (center line), 25th and 75th percentiles (hinges), and 1.5x interquartile range (whiskers).Methods ordered according to their mean Pearson correlation across cell types (rightmost column of Fig. S9B), and cell types ordered according to their maximum Pearson correlation across methods (bottom row of Fig. S9B). Source data are provided as a Source Data file.

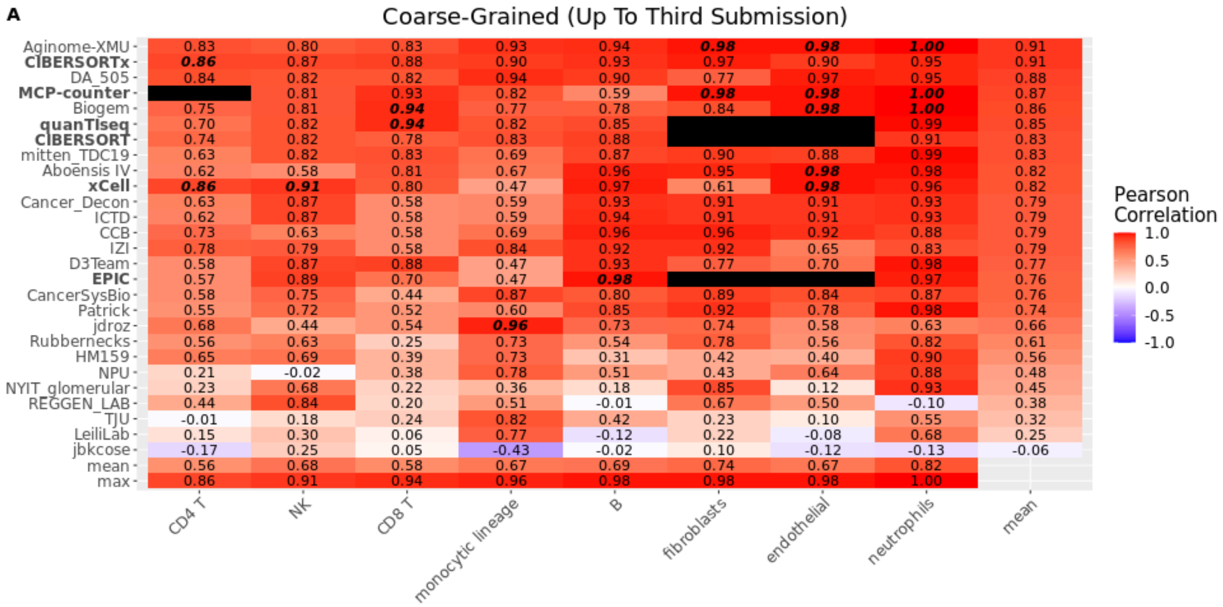


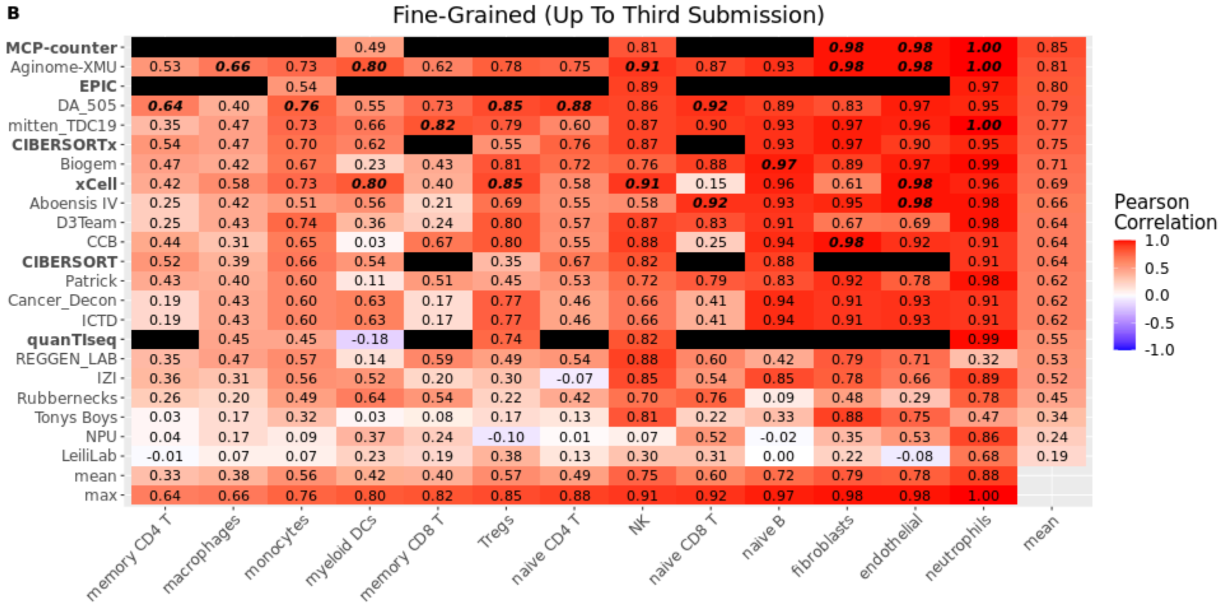


**Fig. S14**: Per-cell type method performance from third submission stratified by sub-Challenge. Pearson correlation of method (left axis) prediction versus known proportion from admixture for each cell type (bottom axis) in (A) coarse- or (B) fine-grained sub-Challenge. Pearson correlation first averaged over validation dataset and then (*n*=1,000) over bootstraps (Methods). Results from latest submission up to third for those methods with fewer than three submissions. Black entry indicates cell type not predicted by corresponding method. Bottom two rows are the mean and maximum correlation, respectively, for corresponding cell type across methods. Rightmost column is mean correlation for corresponding method across predicted cell types. Highest correlation for each cell type highlighted in bold italics. Comparator methods in bold. Methods ordered according to their mean Pearson correlation across cell types (rightmost column), and cell types ordered according to their maximum Pearson correlation across methods (bottom row). Source data are provided as a Source Data file.

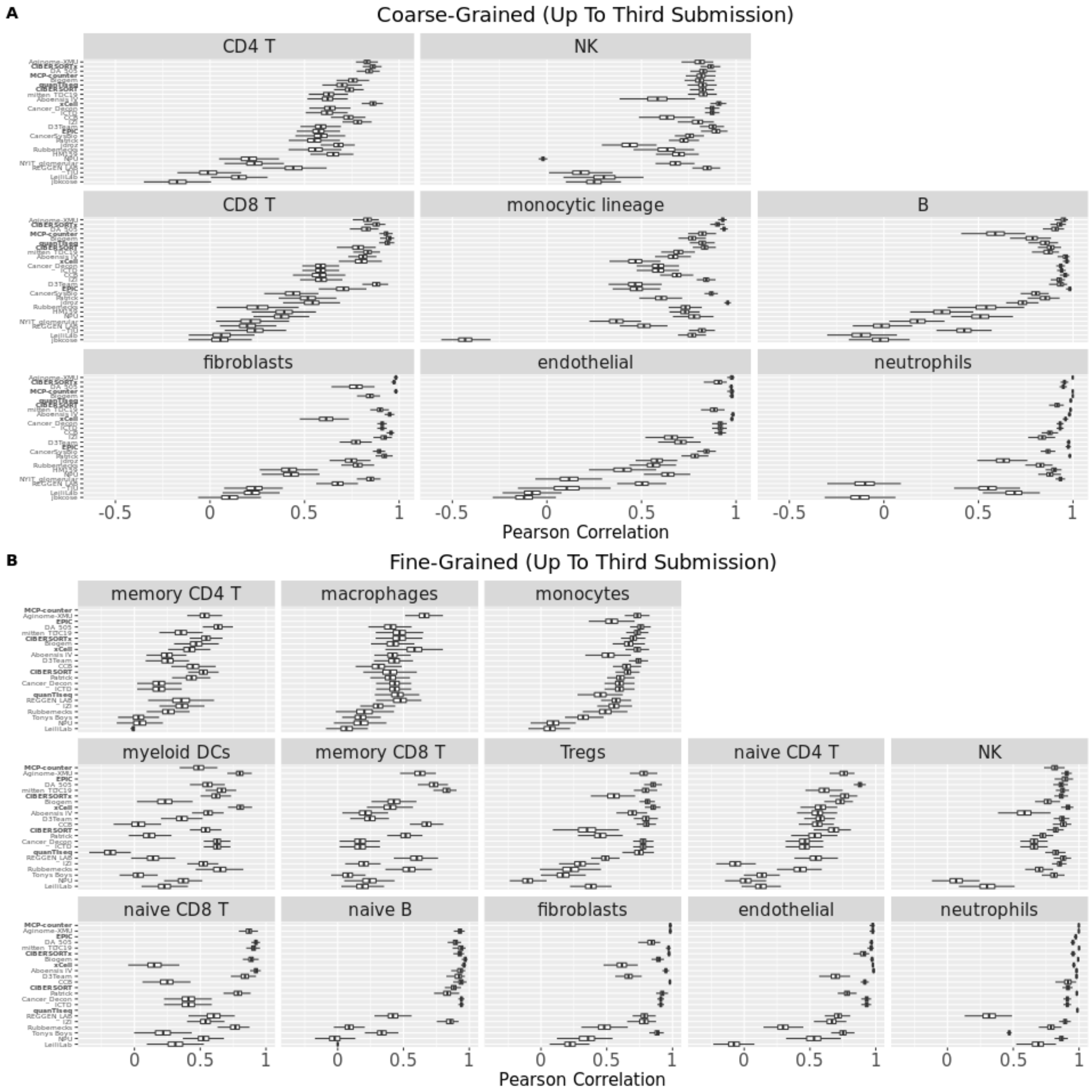


**Fig. S15**: Distribution of per-cell type method performance from third submission stratified by sub-Challenge. Performance (Pearson correlation; x axis) in (A) coarse- and (B) fine-grained sub-Challenges of comparator baseline methods (bold) and participant methods (y axis) for each cell type (facet label). Distribution of Pearson correlations over bootstraps (*n*=1,000; Methods). Results from latest submission up to the third, if less than three submissions for corresponding method. Blank row indicates cell type not reported by the corresponding method. Boxplots display median (center line), 25th and 75th percentiles (hinges), and 1.5x interquartile range (whiskers). Methods ordered according to their mean Pearson correlation across cell types (corresponding rightmost column of Fig. S14), and cell types ordered according to their maximum Pearson correlation across methods (corresponding bottom row of Fig. S14). Source data are provided as a Source Data file.

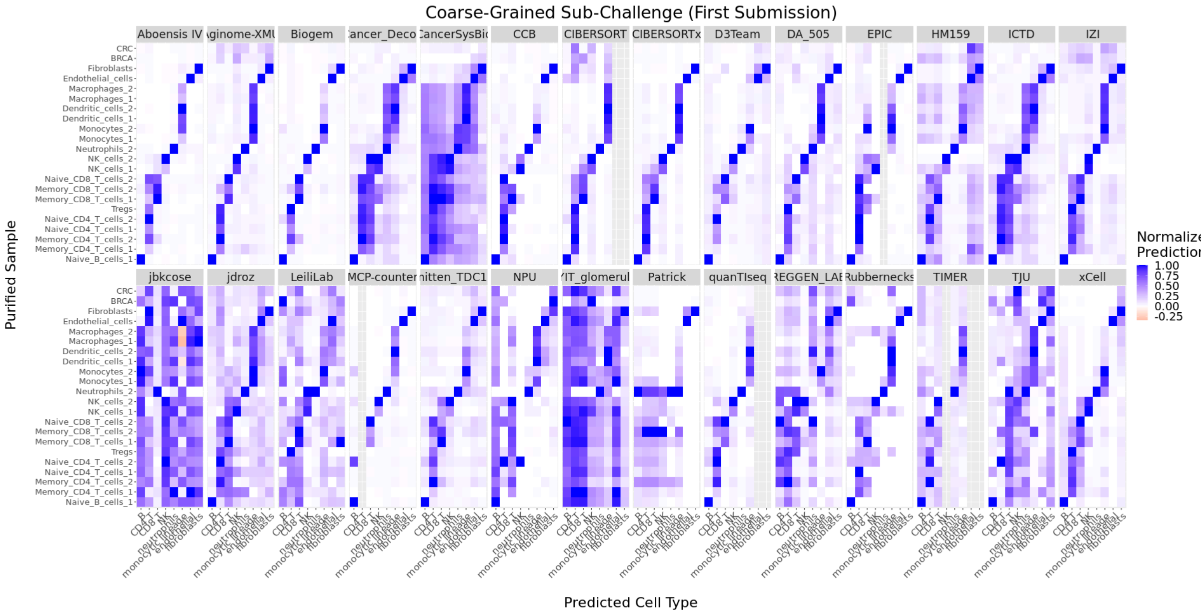


**Fig. S16**: Assessing specificity in coarse-grained sub-Challenge. Normalized prediction of cell type indicated on x axis in purified sample indicated on y axis. Source data are provided as a Source Data file.

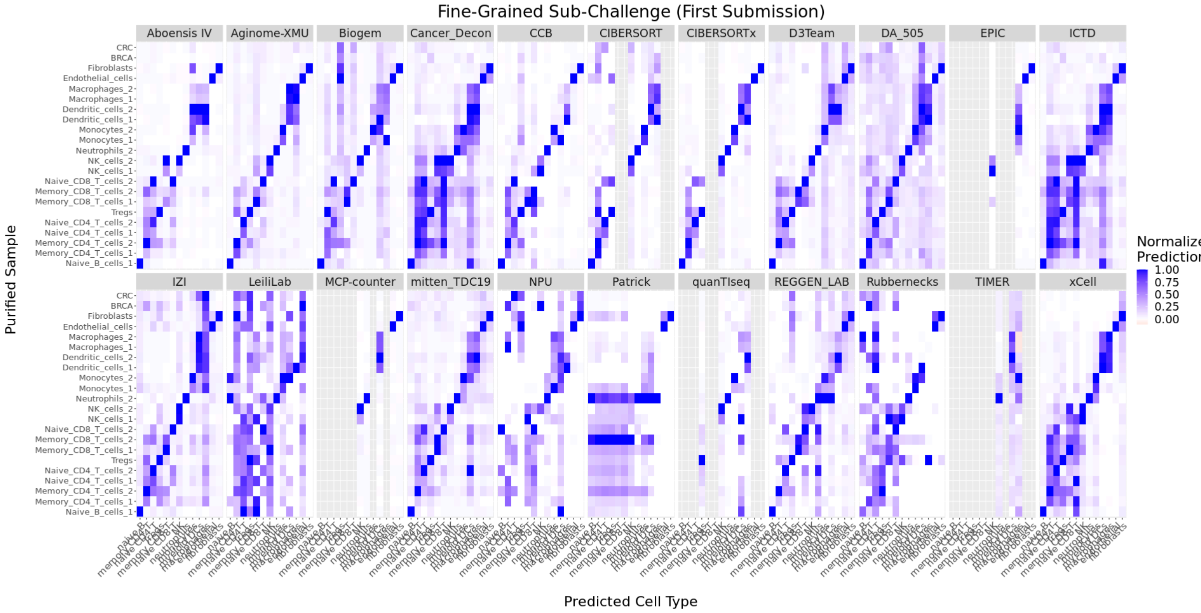


**Fig. S17**: Assessing specificity in fine-grained sub-Challenge. Normalized prediction of cell type indicated on x axis in purified sample indicated on y axis. Source data are provided as a Source Data file.

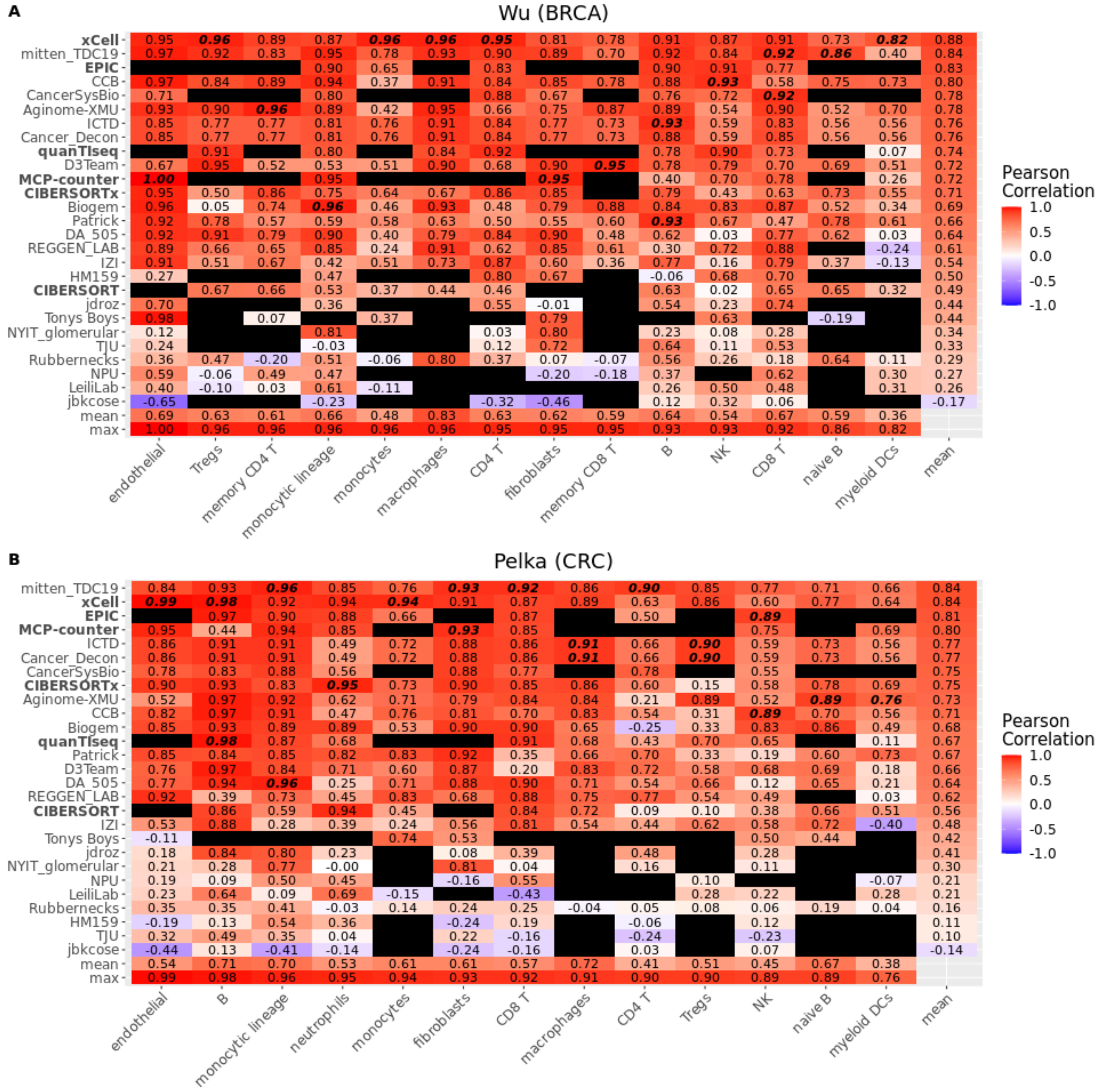


**Fig. S18**: Per-cell type performance of participant and comparator deconvolution methods in simulated cancer datasets. Pearson correlation of method (left axis) prediction versus known proportion from admixture for each cell type (bottom axis) in (A) Wu (BRCA) or (B) Pelka (CRC) datasets. Black entry indicates cell type not predicted by corresponding method. Bottom two rows are the mean and maximum correlation, respectively, for corresponding cell type across methods. Rightmost column is mean correlation for corresponding method across predicted cell types. Highest correlation for each cell type highlighted in bold italics. Comparator methods in bold. Aboensis IV excluded because it failed to predict fine-grained cell types in either dataset. Methods ordered according to their mean Pearson correlation across cell types (rightmost column), and cell types ordered according to their maximum Pearson correlation across methods (bottom row). Source data are provided as a Source Data file.

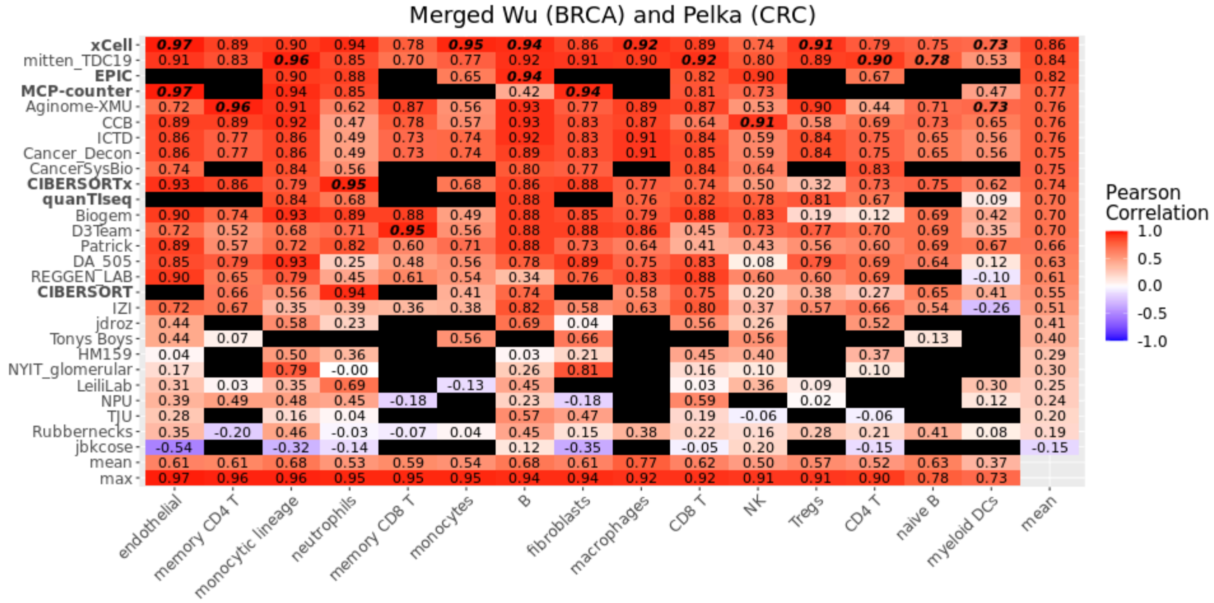


**Fig. S19**: Per-cell type performance of participant and comparator deconvolution methods merged across simulated cancer datasets. Mean pearson correlation of method (left axis) prediction across Wu (BRCA) and Pelka (CRC) datasets versus known proportion from admixture for each cell type (bottom axis). Black entry indicates cell type not predicted by corresponding method. Bottom two rows are the mean and maximum correlation, respectively, for corresponding cell type across methods. Rightmost column is mean correlation for corresponding method across predicted cell types. Highest correlation for each cell type highlighted in bold italics. Comparator methods in bold. Aboensis IV excluded because it failed to predict fine-grained cell types in either dataset. Methods ordered according to their mean Pearson correlation across cell types (rightmost column), and cell types ordered according to their maximum Pearson correlation across methods (bottom row). Source data are provided as a Source Data file.

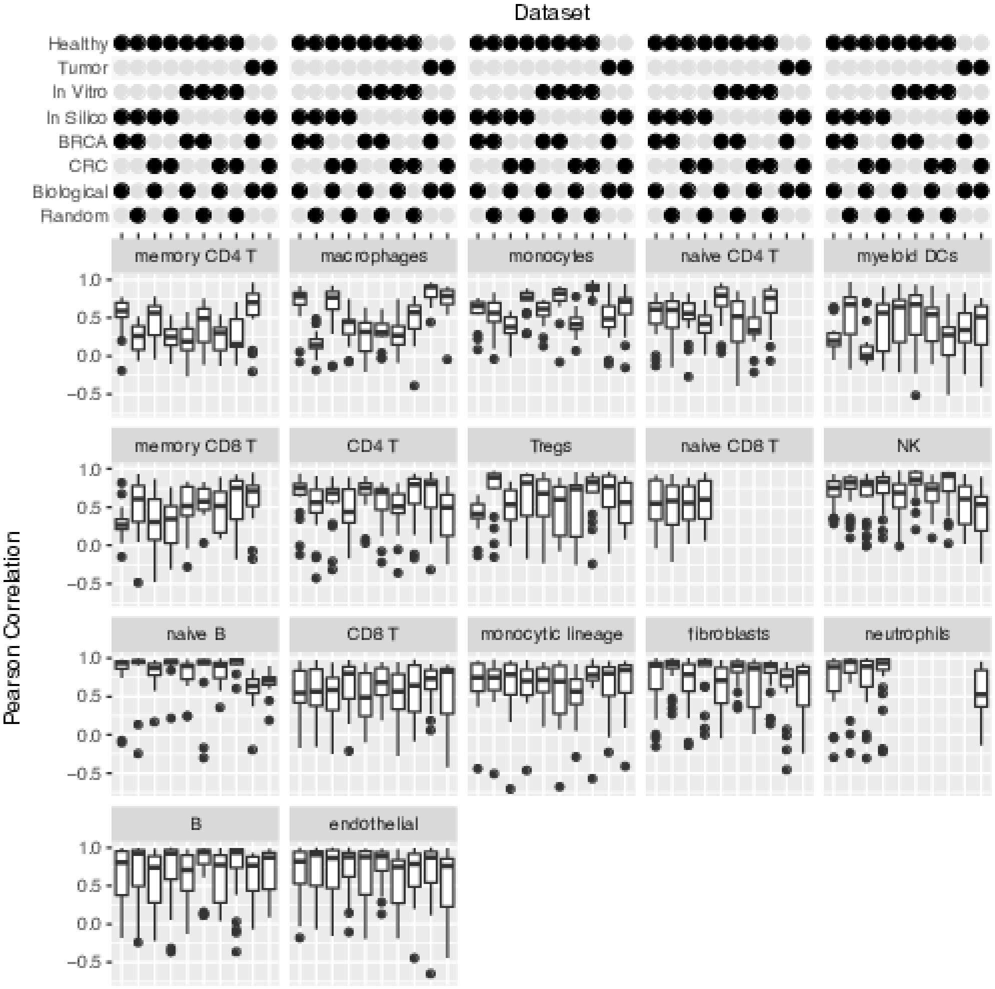


**Fig. S20**: Per-cell type performance of participant and comparator deconvolution methods across datasets with healthy or cancer-associated immune cells. Pearson correlation of method (left axis) prediction versus known proportion from admixture for each cell type across datasets (top axis). Distribution is over all participant and comparator deconvolution methods quantifying the respective cell type. Pearson correlation is averaged over coarse- and fine-grained sub-Challenges for cell types occurring in both. Missing Pearson correlation values indicate cell type absent in the corresponding dataset (e.g., neutrophils in BRCA-associated immune cell dataset). Datasets are characterized based on: 1) inclusion of healthy or tumor-associated immune cells; 2) *in vitro* or *in silico* mixing; 3) inclusion of BRCA or CRC tumor cell expression; and 4) mixing in biologically-informed or random proportions. Aboensis IV excluded because it failed to predict fine-grained cell types in either simulated cancer dataset. Methods ordered according to their mean performance across the ten datasets and the cell types, and cell types ordered according to the max over methods of their mean performance across the three datasets. Source data are provided as a Source Data file.

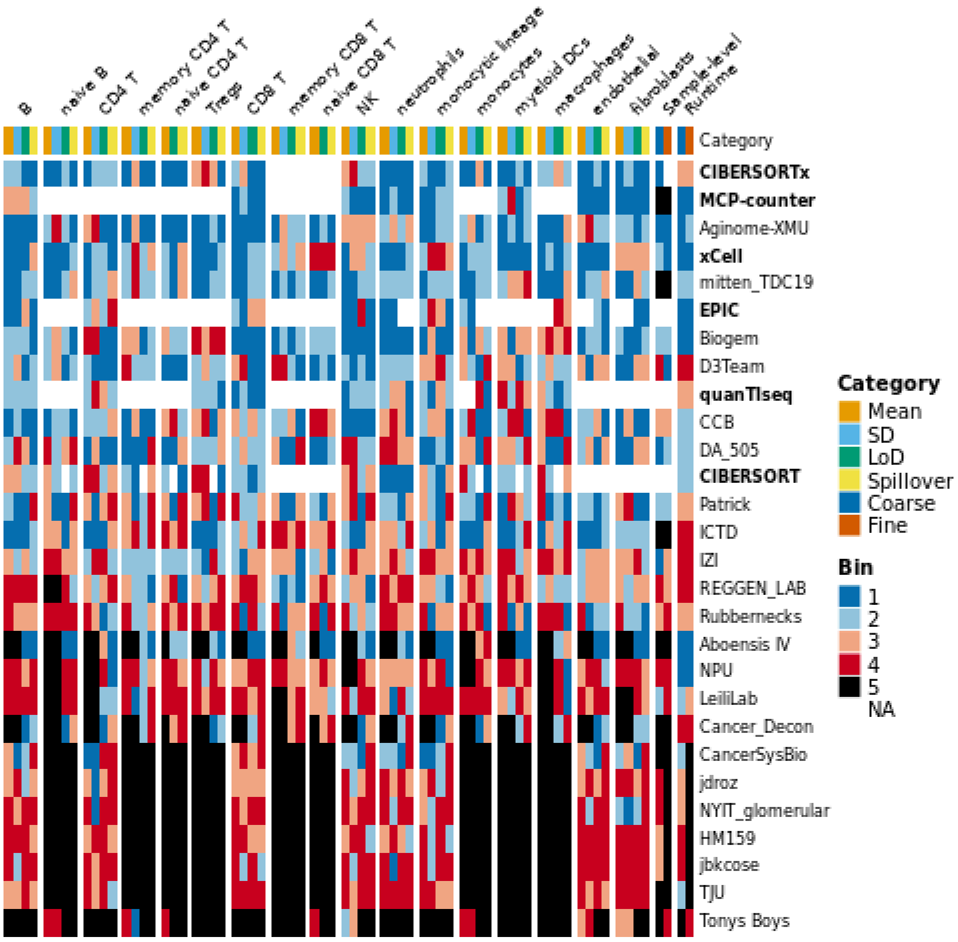


**Fig. S21**: Quantitative summary of participant and comparator deconvolution methods. For each coarse- and fine-grained cell type, methods were assessed across a number of categories: their mean cross-sample, within-cell type Pearson correlation across the Healthy, Pelka (CRC), and Wu (BRCA) datasets, the standard deviation (SD) of their Pearson correlations across those three datasets, and their limit of detection (LoD) and spillover both assessed with the Healthy dataset. Additionally, for both the coarse- and fine-grained sub-Challenges, methods were assessed based on their cross-cell type, within-sample Pearson correlation and their run times. For each category, scores were binned according to quartiles from best (1) to worst (4). Scores that could not be assessed for a comparator method because it was not trained for a particular cell type are marked NA. In all other cases, scores that could not be assessed for a particular participant or comparator method (e.g., because it did not report on the corresponding cell type or was not comparable across cell types) are placed in bin 5. Comparator methods in bold. Methods sorted in descending order according to the mean of their respective bins, excluding NA bins.
